# Neural Implementation of Precise Temporal Patterns in Motor Cortex

**DOI:** 10.1101/2022.04.27.489682

**Authors:** Yuxiao Ning, Tianyu Zheng, Guihua Wan, Jiawei Han, Tengjun Liu, Shaomin Zhang

## Abstract

One of the most concerned problems in neuroscience is how neurons communicate and convey information through spikes. There is abundant evidence in sensory systems to support the use of precise timing of spikes to encode information. However, it remains unknown whether precise temporal patterns could be generated to drive output in the primary motor cortex (M1), a brain area containing ample recurrent connections that may destroy temporal fidelity. Here, we used a novel brain-machine interface that mapped the temporal order and precision of motor cortex activity to the auditory cursor and reward to guide the generation of precise temporal patterns in M1. During the course of learning, rats performed the “temporal neuroprosthetics” in a goal-directed manner with increasing proficiency. Precisely timed spiking activity in M1 was volitionally and robustly produced under this “temporal neuroprosthetics”, demonstrating the feasibility of M1 implementing temporal codes. Population analysis showed that the local network was coordinated in a fine time scale as the overall excitation heightened. Furthermore, we found that the directed connection between neurons assigned to directly control the output (“direct neurons”) was strengthened throughout learning, as well as connections in the subnetwork that contains direct neurons. Network models revealed that excitatory gain and strengthening of subnetwork connectivity transitioned neural states to a more synchronous regime, which improved the sensitivity for coincidence detection and, thus, the precision of spike patterns. Therefore, our results suggested the recurrent connections facilitate the implementation of precise temporal patterns instead of impairing them, which provided new perspectives on the fine-timescale activity and dynamics of M1.

## Introduction

Whether the brain use firing rate code or temporal code has been one of the most fundamental yet controversial question in neuroscience for decades (Abeles (1982); Softky and Koch (1993); Riehle et al. (1997); Shadlen and Newsome (1998); Singer (1999); Decharms and Zador (2000); Brette (2015)). Cracking the neural codes has been considered as the key to understand how information was transformed and processed in the biologically intelligent systems (Barlow et al. (1961); Perkel and Bullock (1968); Rieke et al. (1999); Jazayeri and Movshon (2006)). Apart from its importance in information semantics, neural coding also had implications for neural plasticity and biological learning rules. While spike timing-dependent plasticity (STDP) accentuated the dependency on the temporal relationship between spikes (Bi and Poo (1998); Sjöström et al. (2001)), other forms of Hebbian learning understated it and viewed other variables as more essential than spike timing (Bienenstock et al. (1982); Artola et al. (1990); Ngezahayo et al. (2000); Clopath et al. (2008)).

However, the endeavors to support one view or the other had mostly been made in sensory systems. Motor systems, on the other way, had drawn far less attention on this issue. It was widely accepted that motor systems adapt the rate code to represent movement parameters (Evarts (1968); Cheney and Fetz (1980); Georgopoulos et al. (1982); Riehle and Requin (1989); Moran and Schwartz (1999); Sergio et al. (2005)) or to unfold dynamics for motor preparation and motor control (Churchland et al. (2012); Shenoy et al. (2013); Kaufman et al. (2014); Michaels et al. (2016); Russo et al. (2018)). Moreover, prevalent dimensionality reduction techniques applied in population (Cunningham and Byron (2014); Cunningham and Ghahramani (2015)) and the burgeoning brain-machine interfaces (BMIs) were all built on rate-based models for their computational tractability (Nicolelis (2001); Carmena et al. (2003); Hochberg et al. (2006); Santhanam et al. (2006)). On the contrary, far fewer studies have addressed the implementation of temporal codes in motor systems. The major impediment to sustained vitalization of this area was two-fold. First, researchers typically inherited the representational perspective ingrained in abundant rate-based studies to study the role spike timing played in motor systems. For example, Riehle et al. (1997) had successfully correlated the spike synchronization in motor cortex with the subjects’ expectations of the upcoming “go” cues under a well-designed task structure in the preparatory phase of movement. Similarly, several other evidences for temporal coding in motor cortex had also found covariation between temporal coordination and task-dependent information (Hatsopoulos et al. (1998); Maynard et al. (1999); Baker et al. (2001); Torre et al. (2016)). These studies undoubtedly argued against the autocracy of rate codes in motor cortex, but it was challenging to contrive tasks to reveal the covariation when internal variable rather than overt kinematics variables are involved. In addition, the representational perspective suffered from the lack of “causal power” (Brette (2019)), which could be hard to retake by simply applying manipulation techniques like optogenetics (Baranauskas (2015); Li et al. (2019)) in the cases of studying temporal coding, for they require much higher spatio-temporal resolution to perturb spike timing without hurting the firing structure of uninterested neurons. Thus, a greater understanding of temporal patterns calls for novel paradigms that provide a different perspective from the representational perspective. Second, motor cortex was rich in recurrent connections (Yamawaki and Shepherd (2015)). Recent years have witnessed triumph in fitting M1 activity or testing hypothesis by recurrent neural network (RNN) to reveal the structure or dynamics of M1 that other computational tools failed to capture (Russo et al. (2018); Pandarinath et al. (2018); Ames and Churchland (2019)). This line of studies suggested the recurrent connections might serve for the generation of rich repertoires in M1. However, there are concerns that the recurrent connections might undermine precise temporal structure due to their susceptibility to noise and attractability to chaotic regime. Thus, it seemed unlikely for motor cortex to generate such patterns (Banerjee et al. (2008); London et al. (2010)). Furthermore, following the proposition of synfire chains (Abeles (1991)), some studies showed a correlation between precise timing patterns and the production of song syllables (Chi and Margoliash (2001); Tang et al. (2014)) in the robust nucleus of the arcopallium (RA, vocal motor cortex) that received HVC projections through well-studied synfire chains in songbirds (Albert and Margoliash (1996); Leonardo and Fee (2005); Long et al. (2010)). These pioneering works suggested a rather simple neural structure was indispensable for spikes with precise timing to propagate, also leaving the plausibility and implementation of precise spiking patterns in recurrently connected networks largely unexplored.

Here, we leveraged BMIs to address these challenges. In BMI practices, mappings between neural activity and behavior could be explicitly defined by experimenters. This essentially equipped the researchers with the “causal power” (Golub et al. (2016); Batista (2020)). Some basic neuroscience questions were rejuvenated and reframed by exploiting this “causal power” (Sadtler et al. (2014); Mitani et al. (2018); Hennig et al. (2018); Golub et al. (2018); Clancy and Mrsic-Flogel (2021)). In addition, BMI can be deployed to arbitrary brain areas according to different purposes: S1 in Clancy et al. (2014), V1 in Neely et al. (2018), AIP in Sakellaridi et al. (2019), areas related to the human limbic system and memory in Patel et al. (2021), etc. Hence, by deploying BMI in the primary motor cortex, a highly recurrent brain area that sends commands for movement via spinal cord or downstream brain areas, we could investigate how precise temporal patterns emerged without specified circuit structures. Taken together, we could anticipate the two obstacles mentioned above that harass our further understanding towards the implementation of temporal codes to be overcome by adopting BMI paradigms in motor cortex.

## Results

### Precise temporal pattern in M1 can be volitionally and robustly generated through BMI learning

To investigate whether generating spikes with temporal precision was plausible in highly recurrently connected M1, we developed a novel behavioral task. In this task, the number of concurrent spiking pairs (NCSP) from two randomly selected units (“direct neurons”) in a fixed time window were mapped to the frequency of an audio cursor which was fed back to six rats in real-time. We first estimated the distribution of NCSP using recordings from the baseline block, then set the ~99^th^ percentile of this distribution as the threshold for an artificial reward to guide learning in the following task block (Figure 1A, 1B). if rats up-modulated their NCSP above the threshold within 15s (Figure 1C), the artificial reward was delivered through MFB electrical stimulation (Hernandez et al. (2006)).

**Figure 1.**
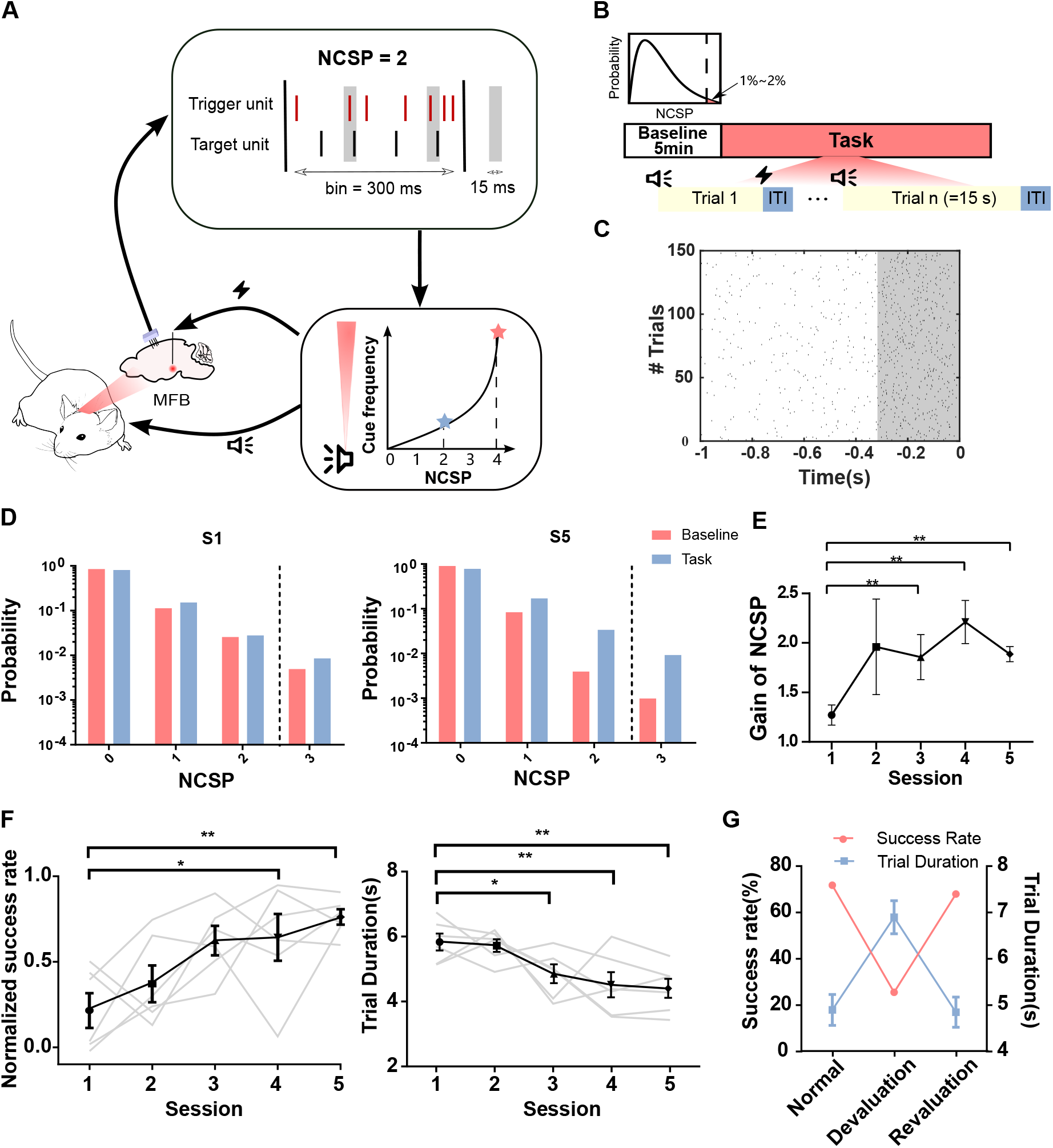
Improvement of performance on a novel behavioral task. (**A**) A closed-loop behavioral paradigm to incentivize volitional and robust modulation of NCSP. In a trial exemplified in the diagram, two concurrent spiking pairs were found from the ongoing neural activity in a 300-ms time bin and mapped to the frequency of auditory feedback as the rats were approaching the goal indicated by the shift from the star in blue to the star in coral. (**B**) A typical session consisted of a 5-minute baseline clock and a task clock. At the end of Baseline, a threshold for rewarding in the ensuing block was set (inset). In the Task, auditory feedback was provided through the trials and was also used as a cue for starting a new trial. Reward was elicited once the rats reached the threshold in 15 seconds, otherwise a failed trial was labeled. (**C**) An raster plot of NCSP aligned to the rewarding time from one session. Each dot represented a concurrent spiking event. It clearly showed a boom of concurrent spiking events in the last bin preceding the reward (shaded area). (**D**) An example of the distribution of NCSP in the first session “S1” and the last session “S5”. The dashed lines indicated the reward thresholds. The Y-axis was set as the logarithmic scale to better reveal the increase in NCSP above the threshold. (**E**) The gain of NCSP increased rapidly and plateaued over the learning course. (**F**) Evolution of task performance. Individual performance was plotted in gray. One-tailed paired t-test, * for p<0.05, ** for p<0.001. (**G**) Manipulation of the contingency between action and outcome.

The marked uplifting of NCSP in the task block appearing in the late phase of learning demonstrated that the rats learned to volitionally modulate NCSP (n = 6, p < 0.01) (Figure 1D, 1E). In addition, the rise of success rate and the shortening of trial duration indicated that rats became proficient at this task in five sessions (Figure 1F). To verify that the causal relationship between the neural activities of chosen units and the outcome was volitionally built under operant conditioning, we performed a devaluation in the last session. The reward was delivered to rats at random times but with a mean delivery rate comparable to the normal block. In the devaluation block, rats were shown to significantly lose their willingness to respond with the drop of success rate and the lengthening of reaction time (Figure 1G). As the contingency was restored in Revaluation, behavioral performance returned to the same level as that in Normal.

The boost of NCSP might arise from grossly increasing firing rates or mediating the precise temporal relationship between the trigger unit and the target unit. Therefore, we used the corrected cross-correlation histogram (CCH) here to reveal the mutual temporal relationship (Gerstein and Perkel (1972); Hatsopoulos et al. (1998); Amarasingham et al. (2012)). We first applied CCH to the 350 ms preceding reward (denoted as “Pre-reward”) in all successful trials of the last session (denoted as “S5”). We found a marked asymmetric peak with short temporal offsets in the corrected CCHs. However, we did not observe such a significant temporal structure in the 350 ms following the start of the trials (denoted “Trial start”) (Figure 2A, 2B). These results indicated that precise temporal coordination contributed to the elevation of NCSP and thus the crossing of the rewarding threshold. Therefore, we reasoned that there should be a session-based refinement of the temporal relationship since the success rates raised and the trial durations dropped from early to late phase of the learning. After applying CCH on all trials in each session (CCH_*task*_), we found the emerging temporal coordination between the paired units when the subjects mastered the task as predicted (Figure 2C, 2D). Besides, the directional preference of the peaks (trigger spikes leading target spikes, not the other way around) found in session-based CCH_*task*_ ruled out the possibility of generally implementing spike synchronization, otherwise symmetrical CCH_*task*_ was expected to be observed which clearly contradicted the result. Therefore, temporally precise neural patterns can be volitionally and robustly generated through BMI learning in M1.

**Figure 2.**
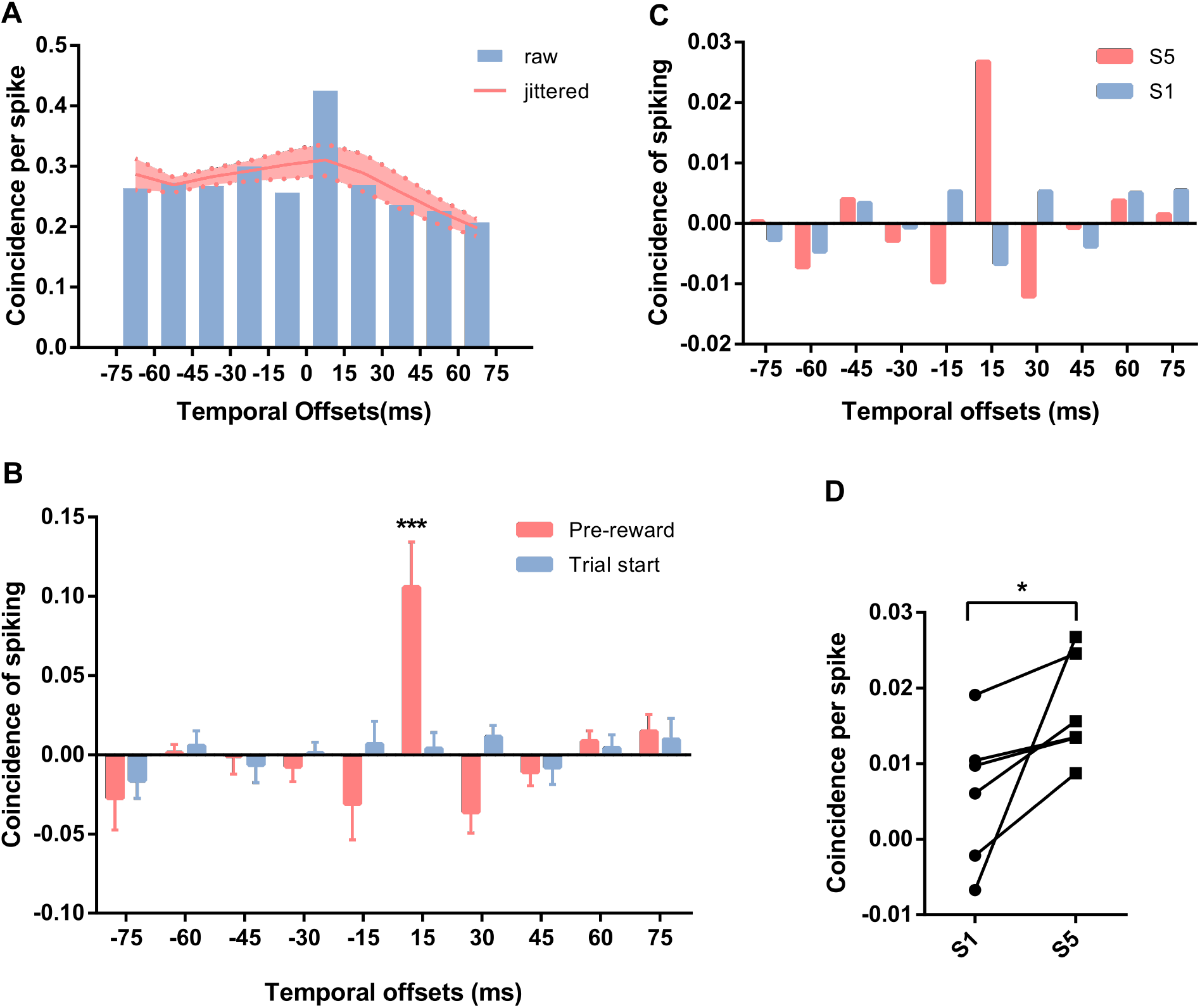
Precise temporal patterns were induced by BMI task with growing rates throughout learning. (**A**) Example raw CCH in Pre-reward of Session 5. The CCH is binned in 15 ms. The coral band represents the chance level inferred by jittering spiking times of the target unit. (**B**) Corrected CCH in Trial start and Pre-reward of Session 5 (n = 6, mean±SD). The coincidence in [0,15] ms is significantly larger than those in other bins for Pre-reward (Repeated measure one-way ANOVA, p = 0.0002; Post-hoc analysis, p < 0.05 for 9 comparisons), while no such precise temporal structure is found in Trial start shown in blue. (**C**) An example of corrected CCH rendered by pooling over all time bins in Task (“CCH_*task*_”) in Session 1 and Session 5. (**D**) Coincidence in [0,15] ms taken from CCH_*task*_ showing the difference between Session 1 and Session 5 (t5 = 2.36, p = 0.032).

Although the cohort exhibited a certain degree of improvement in this particular task, the learning curves and performance at the last session within the cohort were rather at variance. To understand the origin of this variance, we inspected its dependence on the distance between the arbitrarily assigned trigger and target neurons. Converging lines of evidence showed that spike correlation dropped among neurons as the distances increased (Gray et al. (1989); Murthy and Fetz (1996); Berger et al. (2007); Torre et al. (2016)). However, this law only applied partially in our study. For those trigger-target pairs whose distance was below ~600 *μ*m, the monotonic relationship blurred (Figure S1A, S1B). Therefore, we grouped these four subjects as “fast learners”, as they showed better behavioral performance than the two remaining subjects (thus denoted as “poor learners”) (Figure S1C).

### Local network was coordinated at different timescales

Having confirmed the viability for temporally precise neural patterns to be volitionally and robustly generated under the BMI task, we were motivated to investigate how such temporal precision was achieved in a recurrently connected network. Lateral and feedback connections abound in recurrent neural networks, which could mess with the fine temporal structures of neural activities. However, by pruning or inhibiting unnecessary connections through learning, a recurrent neural network could be reduced to execute core computations performed in feedforward structures (DiCarlo et al. (2012); van Bergen and Kriegeskorte (2020)). If this was true in our case, we would observed a precise sequential pattern from neurons in the network (Figure 3A left) which was typically seen in synfire-chain-like structures (Abeles (1991); Diesmann et al. (1999); Ikegaya et al. (2004); Long et al. (2010)). Therefore, we next examined whether the activity of all the neurons recorded (both “direct neurons” and “indirect neurons”) complied with sequentiality with temporal precision. We adapted a handy algorithm proposed in Russo and Durstewitz (2017) to uncover the temporal relationships in neuronal ensembles by detecting cell assemblies on various short timescales (≤ 30 ms), bin widths, and lags. The speculative composition of cell assemblies under the hypothesis of sequential firing should exhibit a spectrum of lags (Figure 3A right). However, though rich types of cell assemblies were found, none of them revealed the synfire-chain-like structures as there was no lag among members in most cell assemblies (Figure 3B). On this account, we rejected the hypothesis that temporal fidelity was carried out by reducing recurrent neural networks to feedforward structures.

**Figure 3.**
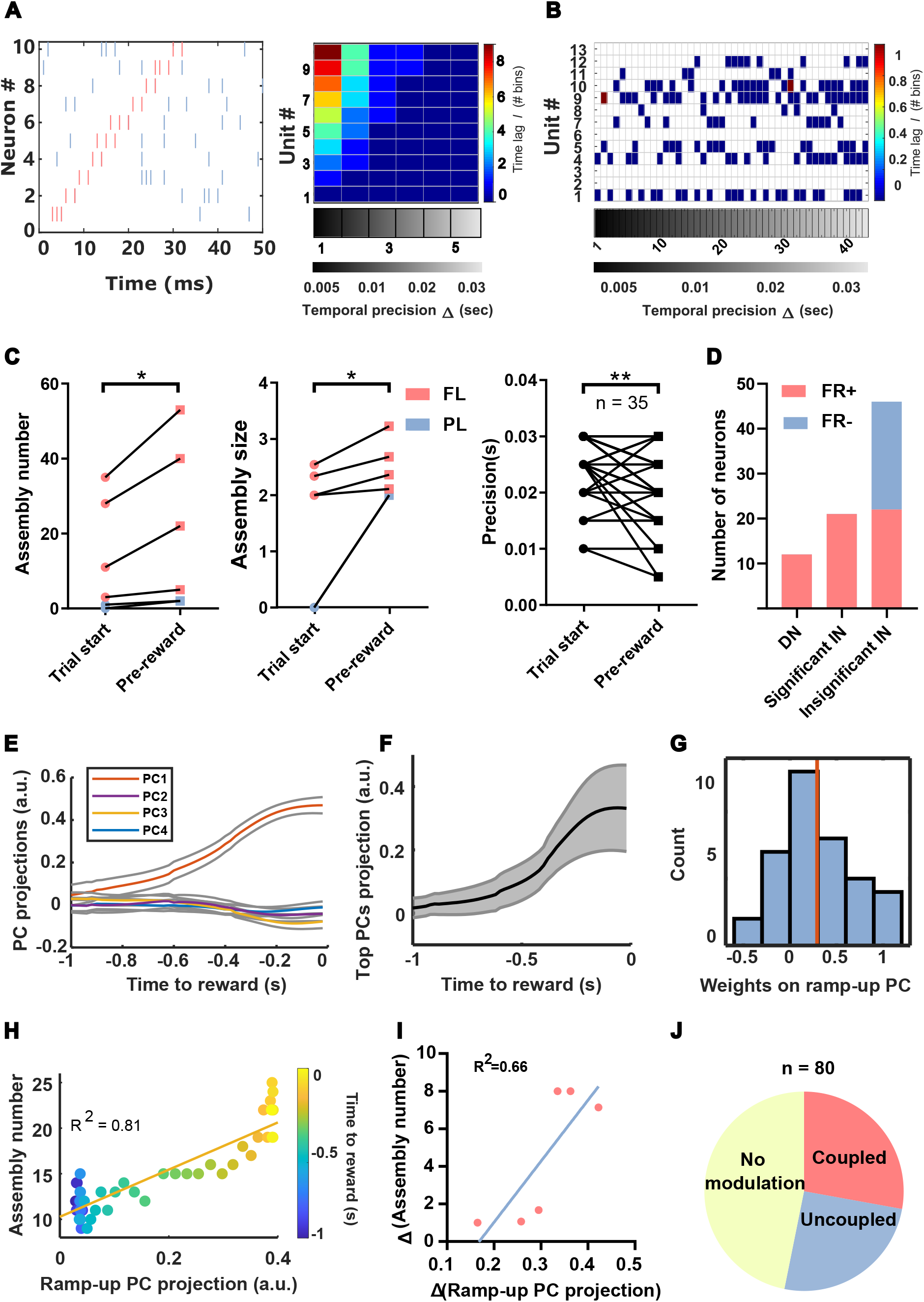
Local network was coordinated in different timescales. (**A**) *left*: Schematic illustrating sequencing firing pattern typically occurred under synfire chains. *right*: Corresponding detected cell assemblies. (**B**) An example showing 44 cell assemblies (columns of the matrix) detected in Pre-reward of the last session. Lags to the leading unit of cell assemblies are color-coded according to the color bar on the right. Cell assemblies are sorted along the abscissa by temporal precision coded by the grey-scale map at the bottom. (**C**) Changes in three assembly-related metrics in session 5. *left*: The number of assembly increased from Trial start to Pre-reward (n = 6 rats, one-tailed Wilcoxon matched-pairs, p = 0.016). *middle*: The size of the assembly increased from Trial start to Pre-reward (n = 6 rats, one-tailed Wilcoxon matched-pairs, p = 0.016). *right*: The precision of assembly increased from Trial start to Pre-reward (n = 35 of cell assemblies, one-tailed Wilcoxon matched-pairs, p = 0.001). (**D**) Mean firing rate changes from Trial start to Pre-reward of three types neurons: direct neurons (“DN”), significantly modulated indirect neurons (“Significant IN”) and insignificantly modulated indirect neurons (“Insignificant IN”) (pooling over all neurons recorded from 6 rats). (**E**) An example (from rat M10) showing temporal projections (1 s preceding reward) on the top 4 principal components ranked by their eigenvalues (mean±SEM). (**F**) Average overall projections of the top 4 PCs by pooling over all rats (n = 6 rats, mean±SEM). (**G**) Histogram of the weights for ramp-up PCs. Mean of weights was greater than zeros, indicated by the red vertical line. (**H**) An example (from rat M10) of correlated increasing of ramp-up PC projection and the number of assembly through time to reward (p = 4.92e-19). (**I**) Correlation between the dynamic range of ramp-up PC projection and that of assembly number (n = 6 rats). (**J**) The proportion of three types of neurons defined by its membership of cell assemblies and firing rates modulation (pooling over all neurons recorded from all 6 rats).

To further understand the implementation of temporally precise pattern in M1, we adopted a bottom-up method by first characterizing the neural population dynamics at different timescales, which then inspired more hypotheses. For relatively short timescales (≤ 30 ms), we continued to use cell assembly for analysis. Cell assemblies were detected in both Trial start and Pre-reward. We found that new cell assemblies emerged as trials proceeded to hit the target and obtain the reward (Figure 3C left). Meanwhile, the assembly size were enlarged (Figure 3C middle). Notably, the number and size of the cell assemblies emerged in “fast learners” was larger than those of “poor learners”. Moreover, the temporal precision of cell assemblies also increased (Figure 3C right). This greater engagement of cell assembly prior to the reward delivery indicated that neural activity was orchestrated in fine timescales. For longer timescales, we first investigated the change of firing rates from Trial start to Pre-reward. Despite of the significant growing of firing rates of all direct neurons, 32.4% of the indirect neurons had significant modulation of their firing rates from Trial start to Pre-reward (one-tailed Wilcoxon matched-pairs test, p < 0.05), among which only one neuron showed declination (1/22). In addition, by comparing the average firing rates of indirect neurons that did not exhibit significant modulation, we found an almost balance number between the increased and the decreased (denoted as “FR+” and “FR-” respectively) (Figure 3D). The single-neuron responses were heterogeneous, yet a net heightening of excitation of the local network was also suggested. To better capture the neural dynamics at the population level, we applied principal components analysis (PCA) to our data by which we could infer continuous neural trajectories as trials proceeded to hit the target and obtain the reward. We observed an overall ramping dynamics from top 3 ~ 4 PCs, which explained 81.3 ± 4.7% of the total variance (Figure 3E, 3F). Further, for those top PCs that exhibit significant elevation (ramp-up PCs, Methods), we confirmed that the majority of neurons exerted positive weights on them (Figure 3G). It was worth noting that 60% of the indirect neurons that showed no modulation in firing rates had positive weights on ramp-up PCs, which implicated the variance stemming from this ramping dynamics was greatly shared through the population. Hence, neurons in the local network were coordinated in both fine and coarse timescales.

Next, we asked if there existed any correlation between dynamics on fine- and coarse-timescales. To draw fine-timescale trajectories, we moving-summed the cell assemblies using a window of 350 ms. We found a co-elevation of the number of cell assemblies and population dynamics in coarse timescales (Figure 3H). In addition, we found the dynamical range of the elevations were also correlated across subjects (Figure 3I). We reasoned that increase of cell assemblies was not likely to be the trivial consequence of increasing global firing rates since the cell assembly detection algorithm have eliminated the contamination brought by variations on coarser timescales. To further confirm this idea, we examined how individual neurons engaged in different timescales. If part of individual neurons exhibited notable temporal relationship with other neurons without increasing its firing rate, we could rule out the possibility that emergence of cell assemblies was directly caused by firing rates modulation.

We thus separated individual neurons into three different groups. If the neuron that exhibited significant modulation in its firing rates was also a member of any cell assembly, it was classified as the “coupled” neuron. Those who did not significantly modulate its firing rates nor were member of any cell assembly were then classified as “no modulation” neurons. The remaining neurons were classified as “uncoupled” neuron, for they showed inconsistent involvement in distinct timescales. Under this classification, we found the population contained all three types of neurons (Figure 3J). In addition, among all “uncoupled” neurons, 20 out of 22 were capable of forming cell assemblies with other “uncoupled” neurons, without the need to engage neurons that increased its firing rates. These results suggested that the raise of excitatory gain revealed through the lens of coarser timescale did not directly contribute to the form of cell assembly at the level of individual neurons.

Therefore, the raise of excitatory gain might facilitate the form and expansion of cell assemblies through network dynamics by transitioning neural regimes, allowing for more neurons to emit spikes with temporal precision. We would took advantage of the network simulation to test this hypothesis in the later sections.

### Functional connectivity was strengthened over the course of learning

Next, we asked what neural structure underlies the learning to generate precise temporal patterns. The causality built by the BMI paradigm endowed researchers with the unique perspective to understand the crucial neural structures supporting or constraining the generation of certain neural patterns (Sadtler et al. (2014); Athalye et al. (2020)). Inheriting this advantage from the BMI paradigm, we could thus differentially characterize the evolution of connectivity between direct neurons and between indirect neurons. We observed a rise in the directed coherence of LFP from the trigger site to the target site through the learning course (Figure 4A up), which suggested the growing functional connectivity from the trigger site to the target site. However, this growing did not hold when reversing the direction from the target site to the trigger site (Figure 4A bottom). We next interrogated the structural changes on network scale by examining all pairwise coherence between the direct sites (where spikes of the direct units were recorded) and indirect sites (where spikes of the indirect units were recorded). Divergent trends in functional connectivity were found at the network level between “fast learners” and “poor learners”, either computed with the trigger site (Figure 4B left, S3A left) or with the target site (Figure 4B right, S3A right). Therefore, although the strengthened functional connectivity from the trigger to the target correlated with the improvement of task performance, it was the strengthened functional connectivity at the network scale underlay the mastering of this “temporal neuroprosthetics”.

**Figure 4.**
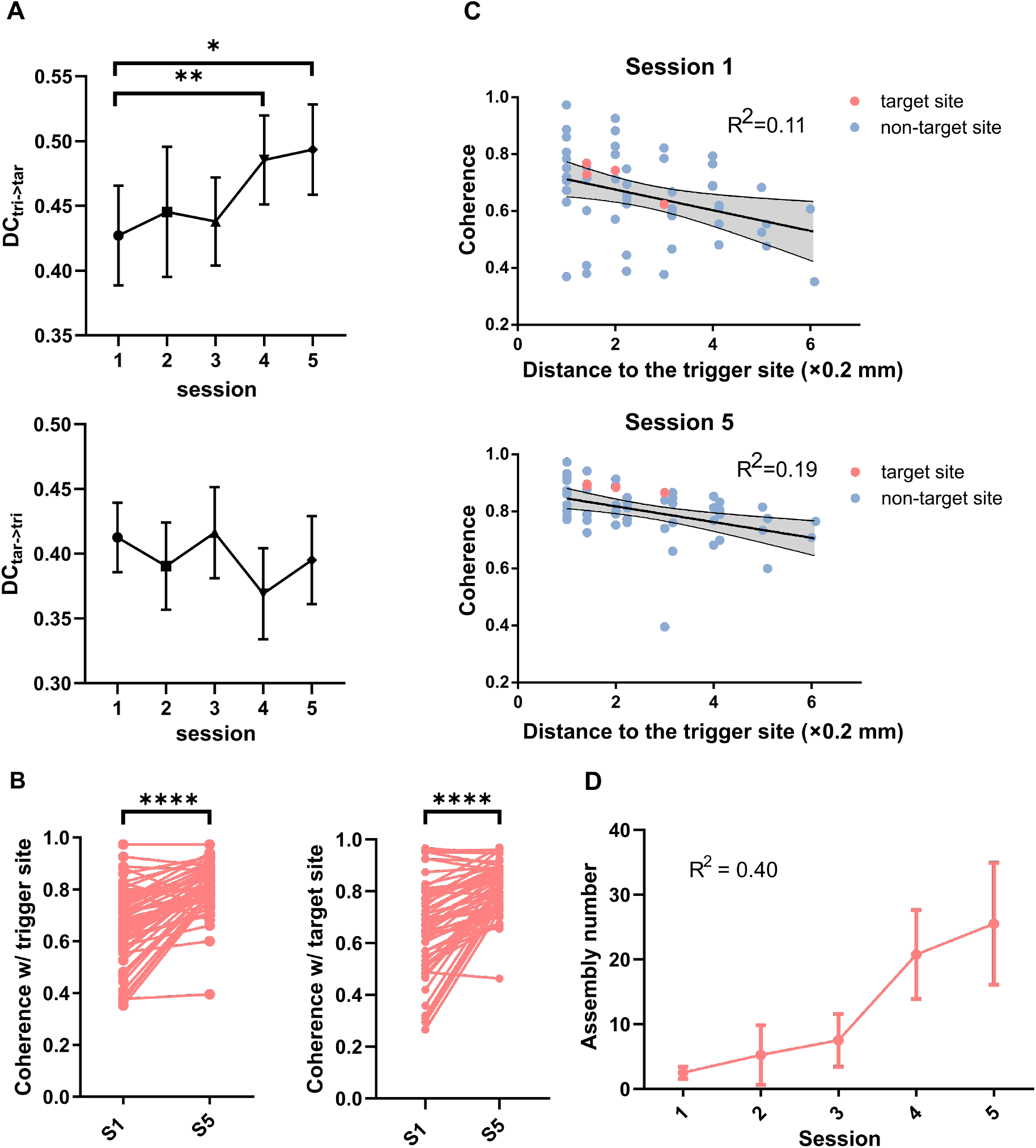
Functional connectivity was strengthened over the course of learning. (**A**) Directed coherence from the trigger site to the target site increased over training (*top*, n = 6 rats, paired t-test: session 4 > session 1, p = 0.004; session 5 > session 1, p = 0.015, mean±SEM). However, no salient trend was found in directed coherence from the target site to the trigger site (*bottom*, mean±SEM). (**B**) *left*: Coherence computed between the trigger site and other indirect sites (paired t-test, n = 56 from 4 “fast learners”, session 5 > session 1, p = 1.03e-11). *right*: Coherence computed between the target site and other indirect sites (paired t-test, n = 56 from 4 “fast learners”, session 5 > session 1, p = 3.19e-11). (**C**) Linear fitting of the distance and coherence of non-target sites to the trigger site (blue, n = 52 from 4 “fast learners”). The shaded band marks the 95% confidence interval. The trigger-target pairs are coded in coral (n = 4 from “fast learners”). *up*: the relation at session 1 (p = 0.013), *bottom*: the relation at session 5 (p = 0.001). **D**. The evolution of assembly number in Pre-reward over the course of learning, error bars represent the SEMs. Linear regression, n = 20 from 4 “fast learners”, p = 0.0026 for slope significantly nonzero.

Despite this gross strengthening of functional connectivity in the network level of “fast learners”, we further asked whether there emerged any more delicate connectivity structure. We depicted the relationship between the spatial distance of paired electrodes and their coherence, and found significant linear correlation at both session 1 (Figure 4C up) and session 5 (Figure 4C bottom). However, the linear relation was more reliable at session 5. Besides, at session 5, the connection between the trigger site and the target site was notably stronger than that of other pairs after being corrected by their distance. The coherence of trigger-target pairs were significantly higher than the 95% upper confidence bound of the linear fitting estimated from all non-target sites. However, the connectivity between direct sites showed no such superiority at session 1.

Additionally, a rather drastic increase in cell assemblies was observed on those “fast learners” (Figure 4D), while only a slight increase was found on “poor learners” (Figure S3B). This divergent trend between the two groups was also seen in the evolution of network connectivity we set forth above. Therefore, we presumed that strengthening functional connectivity within a local network would give rise to the growth of cell assemblies and hence the synchronization of the local network. This hypothesis would be further tested *in silico* in the following sections.

### Excitatory gain transitioned the dynamical regime and facilitated synchronization in the network

We have characterized the neural dynamics through different timescales and found a co-evolution of population excitation and fine temporal structures. However, the elevation of excitation did not correlate with the emergence of fine temporal structure at the single neuron level. Based on these results, we postulated that the elevation of excitation may drive the emergence of fine temporal structures at the network level. Therefore, we sought to intervene in the network with varying degrees of injected excitation to understand its impact on the collective neural activity at fine timescales. To this end, we developed a network model comprising of 4000 excitatory and 1000 inhibitory integrate-and-fire leaky neurons on which the population behaviors could be mediated through two knobs: the external input *ν_ext_* and the ratio of inhibitory postsynaptic potential (IPSP) and EPSP peak amplitudes *g*. Similar to previous studies, mean firing rates, mean coefficient of variation and synchrony were the primary descriptors to characterize the dynamical states of the simulated network (Brunel (2000); Kumar et al. (2008)). The three descriptors could separate the neural space into distinct dynamical regimes, namely synchronous regular (SR), synchronous irregular (SI), and asynchronous irregular (AI) (Figure 5A, 5B, 5C, 5D). Since we found most of the neurons recorded in *vivo* had irregular firing profiles as the coefficient of variation of their interspike intervals (CV_*ISI*_) was greater than 0.7 in both Trial start and Pre-reward periods (Figure S2) (Softky (1993)), we would focus on the analysis in the irregular regime. By marking the synchrony in the irregular regime, we found that as the external input *ν_ext_* increased, the network became more synchronized (Figure 5D, 5E). Interestingly, the gradient of synchrony was larger in the SI regime than that in the AI regime separated by synchrony contour of 0.1, which in term confirmed the separation was a reasonable one as it captured the “phase transition” of synchrony. This transition was also manifested in Figure 5E suggested by the knickpoint. In contrast, as *ν_ext_* increased, the growth of the mean firing rates was rather smooth and linear. Therefore, the elevation of network excitation would transition the neural network to a more synchronous regime. This result explained the previously found emergence of cell assemblies as population activity ramping up (Figure 3H). Besides, the irregular regime requires high *g* indicating a dominant inhibitory effect (Figure 5D), which again argued against synchronization through superfluous correlated excitation exerted on individual neurons and supported synchronization through collective neural dynamics.

**Figure 5.**
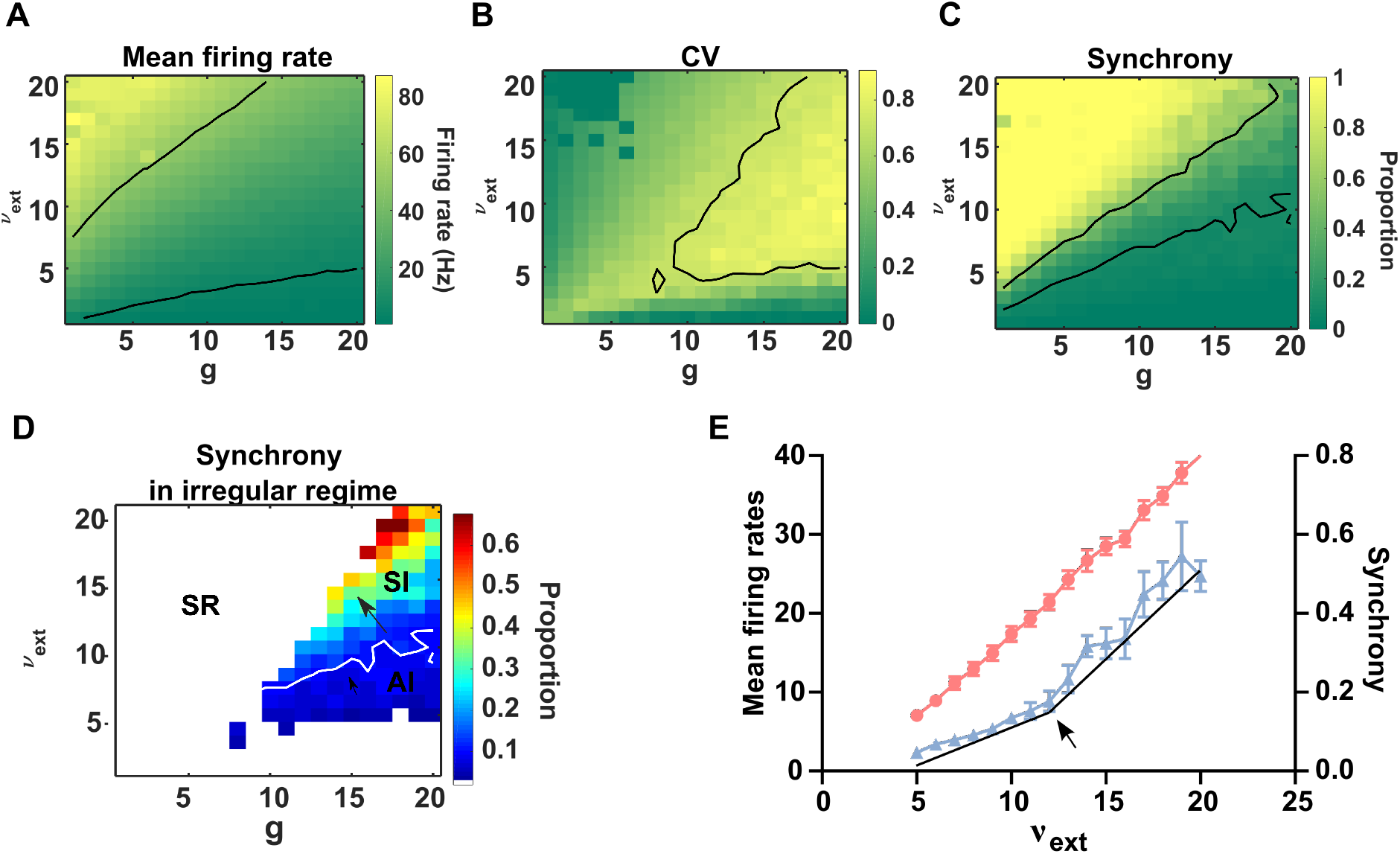
Random balanced network simulation. (**A**) Mean firing rate of the network under parameters *g* and *ν_ext_*. 5 Hz and 50 Hz contours were marked in black. (**B**) Coefficent of variation (CV) under parameters *g* and *ν_ext_*. The contour in the diagram delineated the threshold (0.7) for determining the irregular regime. (**C**) The proportion of significant synchronized pairs to all pairs of neurons in the network (termed as “synchrony”). The contour corresponds to the proportion of 10% and 50%, respectively. (**D**) Gradient synchrony in the irregular regime. The synchrony outside the irregular regime was smeared out. Irregular regime was separated into the synchronous irregular (SI) regime and the asychronous irregular (AI) regime using 10% as the breakline (*white*). Black arrows illustrated the gradient at two regimes. (**E**). In the irregular regime, mean firing rates (*coral*) and synchrony (blue) increased as the external input *ν_ext_* improved, mean ± SEM. *Black* lines represented the slope of the synchrony changes. The knickpoint was marked by the arrow.

### Strengthened connectivity facilitated synchronization and raised subthreshold membrane potential

In Figure 4, we found the network functional connectivity was strengthened throughout learning for “fast learners”. Since the “fast learners” and the “poor learners” showed dissimilar trends in network connectivity and neural population activity in fine timescale, we speculated that this strengthening of network connectivity might play a pivotal role in the generation of global synchronization. Therefore, we need to probe network activity as the network connectivity was strengthened. To characterize the strength of network connectivity, we were inspired by the approach quantifying the clustering in a balanced network proposed by Litwin-Kumar and Doiron (2012). On top of the network model used in the last section, we randomly selected 80 excitatory neurons to form the subnetwork to approximate the underlying network that produced temporally precise dynamics observed from *in vivo* recordings. The degree of functional connectivity of the subnetwork *R_EE_*, was defined as the ratio of two connection probabilities: 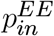 for neuron pairs in the subnetwork and 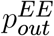 for connections bridging the interior and exterior of the subnetwork. When *R_EE_* = 1, the connections homogeneously distributed in the network, thus no substructures was seen (Figure 6A left). As *R_EE_* increased, salient subnetworks emerged (Figure 6A right).

**Figure 6.**
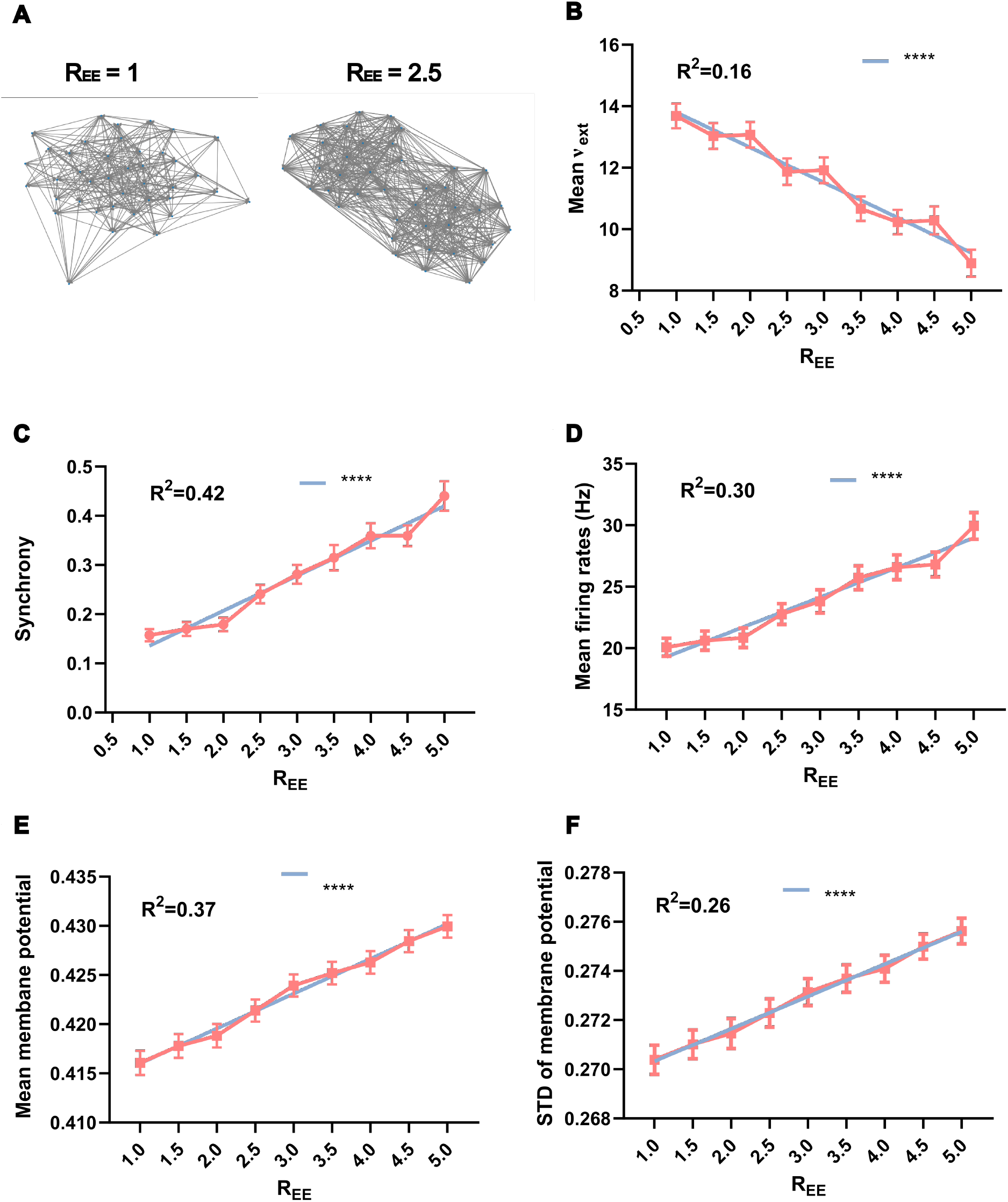
Network after increasing clustering. (**A**) Visualization of networks with different degrees of clustering modulated by *R_EE_*. (**B**) External inputs *v_ext_* of states in the SI regime decreased as *R_EE_* increased (linear fit, n = 574, p < 0.0001 for slope significantly nonzero). (**C**) The synchrony of the overlapped states (overall 27 identical combinations of *v_ext_* and *g* that stably reside in the SI regime under 9 different *R_EE_*) increased as *R_EE_* increased. (**D**) Network mean firing rates at overlapped states in the SI regime increased as *R_EE_* increased. (**F**) The mean membrane potential in the overlapped states in the SI regime increased as *R_EE_* increased. (**C**) ~ (**F**), mean ± SEM, linear fit, n = 243, p < 0.0001 for slope significantly non-zero.

As we turned up *R_EE_*, the mean external stimulus *ν_ext_* needed to drive the subnetwork into the SI state dropped (Figure 6B, S4) while the subnetwork became more synchronous (Figure 6C). Besides, the mean firing rates of the subnetwork increased as *R_EE_* raised given the same *ν_ext_* and *g* (Figure 6D). Together, these results revealed that strengthened connectivity of the subnetwork facilitated synchronization by energy conservation. In addition, we found the subthreshold membrane potential of neurons in the subnetwork raised both in its mean and standard deviation (Figure 6E, 6F).

### BMI learning directed the formation of subnetwork and the generation of reproducible temporal precise patterns

We have successfully applied theories and models of the classic balanced network to elucidated that the elevation of excitatory gain and the strengthening of functional connectivity drive the network to a more synchronous state where average subthreshold membrane potential raised. However, how this population-level dynamics gives rise to the reproducible temporal precise patterns between the assigned direct neurons still remains unclear. From the raised populational membrane potential, we postulated that the target unit might benefit from the raised subthreshold membrane potential to depolarize with higher chance after receiving impulses causally from trigger unit, thus keeping the fidelitous temporal relationship with the trigger unit (Figure 7A). This hypothesis was also able to explain the task learning disparity between “fast learners” and “poor learners”: the strengthened subnetwork connectivity facilitated “fast learners” to enter the synchronous regime with higher subthreshold membrane potential and thus higher discharging precision of direct neurons, while “poor learners” failed to enter the synchronous regime for lacking of structural support. However, since strengthened connectivity between direct neurons was observed in both groups, both groups would exhibit a certain level of improvement in this task. To test this hypothesis, however, we reasoned that the classic balanced networks alone were insufficient as they were mainly used for describing the average and generic dynamics. Specifically, they failed to approximate task-dependant neural activity in three aspects. First, the SI regime simulated from the classic balanced network was chaotic (Hansel and Sompolinsky (1996)) whereas the direct neurons *in vivo* generated rather reproducible temporal precise patterns. Second, the synchronization displayed by classic balanced network was unbiased to the temporal order whereas the direct neurons *in vivo* exhibited precise temporal patterns with notable temporal order (Figure 2C). Third, the original balanced network theory does not involve task-related learning and weights updating whereas the temporal precise patterns was generated through learning. Therefore, we sought to reconcile these discrepancies by introducing modifiable weights and reward-based STDP rule (4) to the classic balanced network.

**Figure 7.**
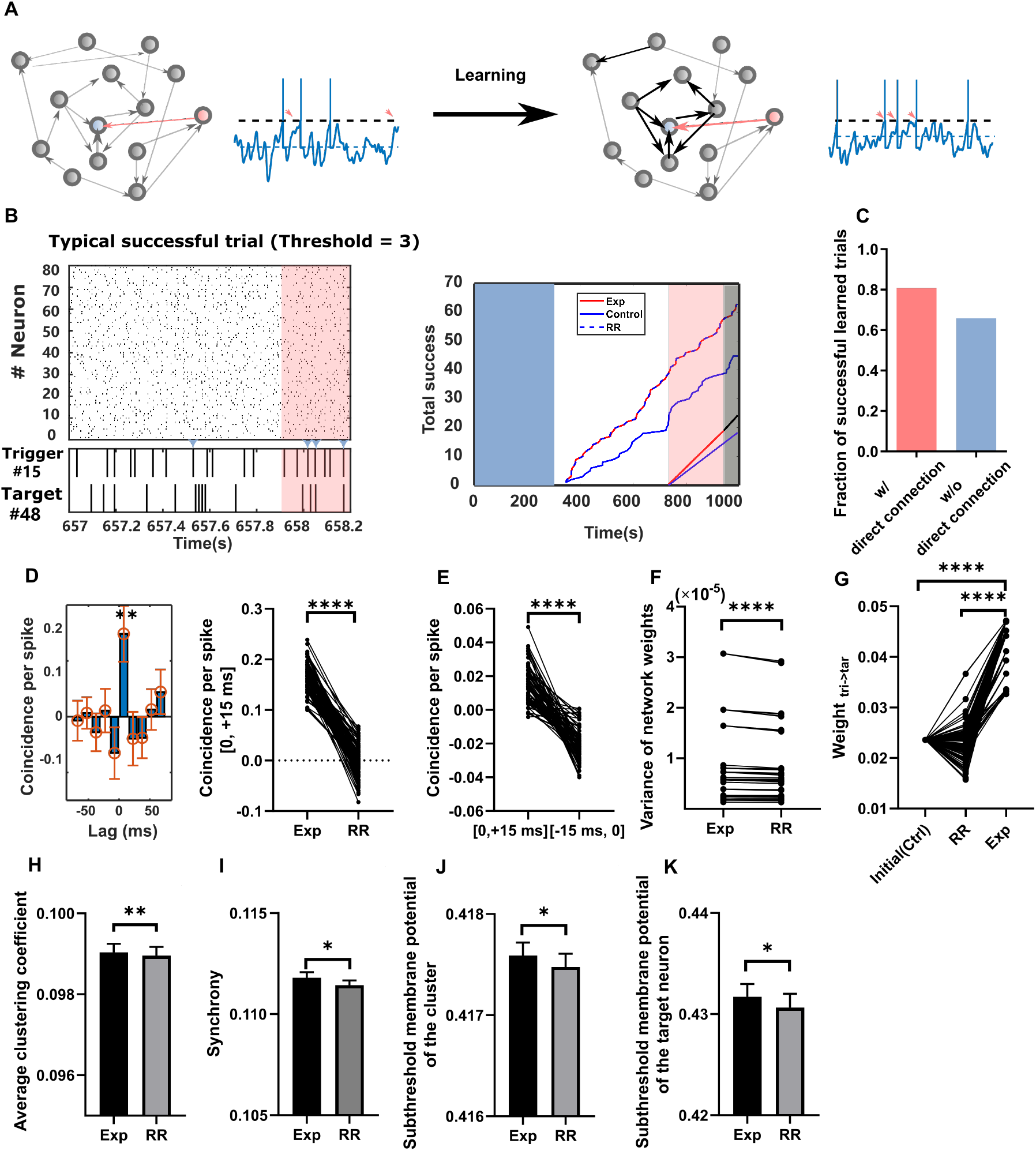
(**A**) Illustration of hypothesized changes on the underlying network and neural activity after learning the “temporal neuroprosthesis”. Trigger neuron and target neuron are color-coded in coral and blue, respectively. Indirect neurons are in gray. The weights of the synaptic connection are coded by the thickness and darkness of the arrows. Membrane potential of the target neuron are shown next to the network model. Angled coral arrows point to the time the trigger neuron spikes. (**B**) *Left*: An example raster plot of 80 excitatory neurons in the network. The activity of trigger neuron and target neuron is zoomed in below. The blue arrows label the concurrent spiking. The shaded coral block marks the last bin of the trial (300 ms) leading reward. *Right*: An example of learning curves in three groups: the experiment group (“Exp”, red), the control group (*blue* solid line) and the random reward group (*blue* dashed line). The blue block marks the 300 s-long baseline. The shaded coral block marks the last 20 bins preceding the termination of the numerical simulation (“Pre-termination”). The following grey block marks a 50 s long simulation where weights were locked for evaluating network synchrony and subthreshold membrane potential after learning (“Weights Locked”). The straight lines below the learning curves indicate the slopes of the learning curves, stacked for comparison. (**C**) Two types of network were trained under the “temporal neuroprosthesis”: the networks with direct connections from the trigger neuron to the target neuron (n = 98 runs of simulation) and the networks without (n = 73 runs of simulation). In majority of simulation runs, the network reached the criterion for successful learning. (**D**) *Left*: The CCH_*pre–reward*_ computed in one exemplified simulation run. The peak appeared in [0, 15] ms bin was consistent with the structure revealed from the *in vivo* experiment. *Right*: Comparison of the coincidence in [0, +15] ms between the experiment group and the random reward group, pooling data from Pre-reward (n = 98, Paired t-test). (**E**) Coincidence in [0, +15] ms and [-15, 0] ms drawn from CCH_*tasfc*_ of the experiment group (n = 98, Paired t-test). (**F**) The variance of trained network weights in the experiment group is larger than the random reward (Wilcoxon matched-pairs signed rank test). (**G**) The directed weights of trigger-target connection under three conditions (n = 98, Paired t-test. Exp > Control, p = 1.32e-58; Exp > RR, p = 1.11e-52). (**H**) The average clustering coefficient of the target-connected subnetwork is higher in the experiment group than the random reward (n = 80 “fast learners”, one-tailed paired t-test, p = 0.006, mean±SEM). (**I**) Mean synchrony of the neurons that projected onto the target neuron is larger in the successful trials (n = 80 “fast learners”, one-tailed paired t-test, p = 0.036, mean±SEM). (**J**) The mean subthreshold membrane potential of the cluster of neurons that project to the target neuron is higher in the experiment group than in the random reward group (n = 80 “fast learners”, one-tailed paired t-test, p = 0.0379, mean±SEM). (**K**) The mean subthreshold membrane potential of the target neuron is higher in the experiment group than in the random reward group (n = 80 “fast learners”, one-tailed paired t-test, p = 0.0319, mean±SEM).

Identical to the *in vivo* BMI paradigm, a pair of neurons was selected randomly from 400 excitatory neurons as directed neurons whose NCSP was used for rewarding and thus updating the weights. Meanwhile, the parameters were set to assure that the network was operated in SI regime. In this way, we could probe whether reproducible and asymmetrical temporal patterns between direct neurons could be evolved from the chaotic and symmetrical synchronized plain network while complying with the raised subthreshold membrane potential predicted by the hypothesis.

Besides the simulation on plastic network (denoted as “Exp”), we also ran two types of controlled simulation. In the random reward group (“RR”), the reward signals occurred within a ±3 s jitter window to the rewards in Exp. In the control group (“Control”), the weights were fixed as time elapsed. We found that in most simulation trials, plastic networks learned to generate desired neural patterns between direct neurons with both reproducibility and temporal order (Figure 7B left). The networks in Exp outperformed the plain networks (“Control”) (Figure 7B right). The learnability was obvious even if no direct connection existed between the trigger and target units, albeit slightly lower than networks containing direct connections (Figure 7C). To confirm the network was trained to quasi-optimal while we terminated the weights updating, we locked the weights and continuously tracked the performance for another 50 s to examine the stability of success rate. The success counts per second estimated in Pre-termination (time spanning last 20 success) and in Weights Locked (50 s after simulation was terminated) were higher than that of Control, assuring that network was quasi-optimal by the time we terminated the training (Supplementary Figure S5A). The small portion of networks that failed to meet the criteria for termination in the maximum learning time also exhibited certain level of improvement (Supplementary Figure S5B). Therefore, we referred this group of networks as “poor learners” who showed similar deficiency in learning the “temporal neuroprosthetics” as those “poor learners” in the in vivo experiment. In parallel, the remaining networks was referred as “fast learners” in the simulation.

We also depicted the CCH of neural activity in the Pre-reward to exam the temporal relationship between direct neurons (Figure 7D left). The peak that appeared at [0, +15] ms was consistent with the *in vivo* findings. The salient contrast to the coincidence drew from RR group further demonstrated that the reward-based network was capable of learning to generate temporal precise patterns with reproducibiliy (Figure 7D right). By depicting CCH from Pre-termination, we confirmed the temporal precise patterns generated were asymmetrical (Figure 7E). Therefore, we verified that the generation of reproducible and asymmetrical temporal patterns with precision was learnable under reward-based synaptic plasticity on top of the SI regime in balanced networks.

After learning, the weights became heterogeneous compared to the uniform initial weights (Figure 7F). Besides, beyond the heterogeneity brought by unbiased rewarding revealed by the RR group, some fine structure emerged in Exp which might support task-related improvement (Figure S5C). This motivated us to seek the network structure underlying the learning of “temporal neuroprosthetics”. We first interrogated the connection from trigger neuron to target neuron, and found its weight increased asymptotically to the maximum weight *W_max_* (Figure 7G, S5D). This was in line with the increased directed connectivity in both “fast learners” and “poor learners” shown in Figure 4A. We next investigated the whether there emerged or strengthened some subnetwork by computing the average clustering coefficient (Methods) of set of neurons that connected to the target neuron. The average clustering coefficient of “fast learners” was higher than that of the corresponding random reward group (Figure 7H), suggesting the formation and / or strengthening of the subnetwork containing the target neuron. This result also aligned with the strengthening of network connectivity revealed in Figure 4B. Moreover, the “poor learners” did not show growth in the average clustering coefficient which was significantly lower than that of the “fast learners” (Figure S5E). According to the hypothesis, strengthened network connectivity facilitated entering to a more synchronous state where the subthreshold membrane potential of the subnetwork and the target neuron inside should be raised. Indeed, we found the synchrony (Figure 7I), the subthreshold membrane potential of the subnetwork (Figure 7J) and the subthreshold membrane potential of target neuron (Figure 7K) all increased compared to the random reward group without modulation in average firing rates (Figure S5F), which supported our hypothesis: the form and / or strengthening of recurrent subnetwork improved the sensitivity for coincidence detection (Figure 7A). Moreover, they all showed disparity between “fast leaners” and “poor learners” (Figure S5G, S5H, S5I), which suggested that the strengthening of network connectivity also accounted for the differences in learning rate.

## Discussion

To our knowledge, this work is the first to use operant conditioning to robustly recruit precise spiking on a millisecond time scale with causal relationship with outcomes. One can thus be freed from laboriously searching for the statistical relationship between temporal patterns and “decoded” parameters which may be confounding, especially when internal variables are involved. Besides, leveraging the BMI paradigm provides one more insight into neural implementation of certain patterns or algorithms (generation of precise temporal patterns in our study) as reveals biological constraints (Sadtler et al. (2014); Clancy and Mrsic-Flogel (2021)). In addition, we are the few to pin down the “rate-based vs. spike-based” argument to the primary motor cortex (M1). Three potential challenges seemingly made M1 a less ideal place for generating spike-base codes: i). M1 mainly receives inputs from other cortical areas and thalamus and forms abundant strong reciprocal connections with them (Muñoz-Castañeda et al. (2021)). Meanwhile, there are massive intrinsic recurrent connections in M1(**?**). This highly recurrent environment can destroy the temporal fidelity between spikes; ii). M1 is absent of clear topographic structure compared to primary sensory cortices, which have long been the main battlefield for the neural coding debate (Hatsopoulos (2010)). This also adds up complexity; iii). M1 sends its projection directly to spinal interneurons and motor units. However, muscle reaction times are often hundreds of milliseconds long, which is way longer than a complete spike or spike timing offsets that are conventionally regarded as precise. Therefore, the temporal resolution of M1 may be constrained by the biomechanics of the effectors. Given the above rationales for underestimating the motor cortex to produce precise spike patterns, our results yet extended spike-based theories to motor cortex.

One might argue that the precise temporal patterns were generated under abstract skills in our study, not from the natural repertoire, thus the results and implications gained from our study were trivial for understanding how natural behaviors are produced. However, we disapprove of this view in two ways. First, unlike most BMI tasks that strictly prohibit overt movement, we did not prevent subjects from freely moving. It was shown in **??**(Supp movie) that the subject indeed made overt movements while up-modulating its NCSP to obtain rewards. On the contrary, Chapin et al. (1999) have found that the animals were prone to the ’neurorobotic’ mode and reduced physical movement once they learned to use neural signals to manipulate the robot arm and obtain reward. From this we can infer, the precise temporal patterns produced in our experiment were not detachable from endogenously associative movements. Therefore, the underlying neural activity for natural behaviors should be partially shared to generate the precise temporal patterns under BMI paradigm. Second, the BMI task provided insight into the biological constraints that were usually difficult to obtain following traditional behavioral tasks (Sadtler et al. (2014); Athalye et al. (2020)). In this study, for example, by varying the distance between the trigger unit and target unit, we found that the distance was a limiting factor to generating precise temporal patterns. These biological constraints can inspire more biologically plausible hypothesis and models for generation of natural behaviors.

The results of the experiment in vivo showed the coevolution of cell assemblies and network excitation both in single-trials (Figure 3H) and cross sessions (Figure 4D). However, this covariation did not apply to all individual neurons (Figure 3J). Thus, we postulated that excitatory gain of the network indirectly caused the emerging and expansion of cell assemblies by driving the neural states to a more synchronized regime. This idea had been supported by the numerical simulation (Figure 5E). The degree of synchrony rest on the energy injected into the network. Because the degree of synchrony in the numerical simulation was a fair approximation of the number of cell assemblies in vivo, we validated that the emerging and extension of the cell assemblies depend on the energy injected into the network. Noted that the neurons in this regime are subject to fast and strong inhibition, higher injected energy paves the road for recruiting more concerted spikings. If we adopt the language of dynamical systems, the SI regime was an “attractor”. In addition, from the perspective of dynamical systems, strengthening the network connection (Figure 6B, 6D) meant deepening the potential well in the energy landscape, adding more attractiveness so that less external energy would be required to keep the neural activity in SI and fuel more cell assemblies (Figure 6C) (Hopfield (1982); Litwin-Kumar and Doiron (2012)). This might explain why we had observed those rich cell assemblies in the late phase of learning (Figure 4D).

Spike-based theories, as well as the possibility of robustly producing spiking patterns, have been questioned based on the variability of spike trains observed both in trials and over trials (“trial-to-trial variability”) (Shadlen and Newsome (1998)). This unreliability was often modeled as intrinsic noise subjected to Poisson distribution. However, if relative timing was taken into account, we demonstrated that NCSP could be modulated with frequency and temporal precision well beyond the chance level (Figure 1D). In other words, the exact neural trajectories and spike timings might be subject to initial conditions or perturbations (Banerjee et al. (2008)), but population temporal interactions could be reserved.

It is worth noting that in the study of Engelhard et al. (2013), precise spike synchrony in M1 was also induced by adopting BMI. However, the specific patterns and underlying mechanisms in their practice are different from ours. The hallmarks accompanying the precise spike synchrony in their study: enhanced power of gamma oscillation and stronger phase locking were not seen in our study (Figure S6A, B). In fact, oscillation has long been implicated as an “internal clock” to pattern the neural activity in temporal order(Treisman (1963); Lubenov and Siapas (2009)). Therefore, we also examine whether there was significant modulation of theta band power or phase locking, which was widely seen in the hippocampus for encoding spatial information with precise timed patterns (Buzsáki (2002); Huxter et al. (2003)). Nevertheless, we did not find power change or phase locking in theta band of our data (Figure S6C, D). Thus, neural oscillations are not necessarily a scaffold to frame spike timing to produce precise temporal patterns.

We should clarify that brain areas underlying the generation of precise temporal patterns in M1 are not necessarily constrained in M1. Similarly, plasticity that leads to the strengthening of functional connectivity in the subnetwork may also occur in multiple locations. This is the classical credit assignment problem. We proposed that the actual modification of synaptic connectivity might happen in i). M1, as dopaminergic projections to M1 were found to mediate motor skill learning (Hosp et al. (2011); Guo et al. (2015)). Besides, there are lines of studies showing that synaptic connectivity in M1 is susceptible to change after motor learning (Rioult-Pedotti et al. (1998); Kleim et al. (2004) Hayashi-Takagi et al. (2015)). Therefore, changes in motor cortical structural connectivity are seen as the basis for long-term motor memory; ii). Other cortical or subcortical areas that have projections to M1 (and those projections), such as the premotor cortex (Perich et al. (2018)), thalamus(Biane et al. (2016); Hasegawa et al. (2020)) or basal ganglia through the cortico-basal ganglia-thalamo-cortical loop (Shen et al. (2008)); iii). The combination of all the above candidates. The synaptic changes at these candidate locations can be evoked in parallel or in series (Kupferschmidt et al. (2017)). We did not disambiguate these possibilities in our analysis. Instead, we have abstracted all the related mediations of plasticity into the dynamics of functional connectivity. In this way, we can bypass the credit assignment conundrum to focus on understanding the how strengthening population connectivity facilitate the generation of precise temporal patterns.

One of the exciting findings in our study is that the functional connectivity between arbitrarily assigned two sites was strengthened. This was first revealed in our study by the cross-correlation histogram, which can be used to characterize functional connectivity despite its wide use in uncovering temporal relations (Brody (1999); Denman and Contreras (2014); Kobayashi et al. (2019)). Additionally, we used LFP signals to infer functional connectivity to avoid the needle-in-the-haystack problem brought by spike-based inferences. Model-based directed coherence and model-free coherence were applied (Srinivasan et al. (2007)) to reveal the strengthening of connectivity over learning. Notably, the connectivity between the assigned sites by experimenter was prominently higher than other connections, which indicates potential of applying our “temporal neuroprosthetics” to guide the structure of connectivity (Figure 4C). It can be a step forward in engineering plasticity (Jackson et al. (2006); Moritz (2018)) and rehabilitation from neural diseases caused by loss of neural connections. Furthermore, inducing plasticity by operant conditioning is a natural way of learning that does not require the introduction of any artificial stimulation into the brain. However, plasticity engineering in this way becomes challenging when the distance between two inducing sites is larger than some critical values, as shown by the performance of “poor learners” in our study. Although directed connectivity from the trigger site to the target site was also strengthened, the background network collapsed to provide additional support seen in ’fast learners’ (Figure S3A). We assumed that this constraint might arise from: i). Innately lower connection probability and sparser connections in the background covering two sites that are farther apart; ii). Inhibitory mechanisms that prevent co-firing in a rather large area, which may lead to pathological discharging. Future studies are needed to test these hypotheses.

The strengthening of functional connectivity between direct neurons may be accelerated by well-designed BMI “decoders”. The temporal criterion for concurrent spiking pairs (<15ms) was in the synaptic potentiation window according to the STDP rules. If exists, synapse from the trigger neuron to the target neuron would be expected to be potentiated by reward-based STDP. Furthermore, the task encourages the production of concurrent spiking pairs at a high rate (higher than 98% of the time). This is equivalent to conducting more pairing trials for potentiation, which in turn facilitates generating more concurrent spiking pairs.

Our study suggested that external input in a certain range is critical to drive the neural state to a SI regime, which set the foundation for generating neural patterns that resemble realistic firing profiles. This inevitably caused more spikes to be emitted in the network (Figure 6D). However, one of the advantages of temporal code that has been frequently suggested is its energy efficiency (Olshausen and Field (2004); Rehn and Sommer (2007); Jadhav et al. (2009); Stöckl and Maass (2021)). It was also an appealing feature of spiking neural networks (SNN) and neuromorphic computing. (Davies et al. (2018); Pfeiffer and Pfeil (2018); Taherkhani et al. (2020)). Thus, this gap needs to be filled to deepen our understanding of the temporal coding issue. Modifying the present BMI “decoder” to constrain the level of excitation may be a favorable solution in future studies.

## Methods

### Surgical procedures

All surgical and experimental procedures conformed to the Guide for the Care and Use of Laboratory Animals (China Ministry of Health) and were approved by the Animal Care Committee of Zhejiang University, China.

Six male Sprague-Dawley rats weighing approximately 250 grams were used in the experiments. After anesthesia (Propofol 10 mg/kg, 10 ml/kg body weight), each rat was placed in a stereotaxic apparatus and chronically implanted with a 16 channel nichrome microwire array (2×8 configuration, 35 *μ*m diameter, 200 *μ*m electrode spacing) in primary motor cortex (AP: +2.3 mm; ML: +2.5 mm; DV: +1.25 mm) and stimulating electrodes made from pairs of insulated nichrome wires (65 *μ*m diameter) with a 0.5 1mm vertical tip separation in medial forebrain bundle (MFB, AP: −3.6 mm; ML: +1.8 mm; DV: +8.7 mm) bilaterally. Rats were given 7-8 days to recover after surgery before behavioral training.

### Behavioral task

The activity of the trigger and target units was binned every 300 ms. The action potentials of the target unit following those of the trigger unit in a short delay (< 15 ms) were defined as a concurrent spiking pair. Each action potential in a bin was included in no more than one concurrent spiking pair. The number of concurrent spiking in each bin was detected and exponentially mapped to the pitch of an audio cursor ranging from 1-24 kHz in a quarter-octave increment (Koralek et al. (2012)) by the custom-written online program in Matlab (Mathworks, USA). This feedback signal was further smoothed with moving averages in three consecutive control bins. Rats moved the cursor to higher pitches by up-modulating the number of concurrent spiking pairs until the goal was reached within 15 seconds. Otherwise, a failed trial was labeled. MFB stimulation was delivered as long as the target was hit. The 4 s long intertrial interval was set to eliminate potential interference brought by electrical stimulation and prepare the rats for the next trial. During the baseline block in each session, the number of concurrent spiking pairs (NCSP) was sampled every 300 ms (to yield 1000 samples in 5 minutes). We then draw a histogram to approximate the distribution of NCSP, based on which the rewarding thresholds were set. Specifically, we follow two principles to set the threshold: i). The corresponding percentile of NCSP should be around 99%, no lesser than 98% to guarantee that target achievement in a 15 seconds long trial was a small probability event if no effort was paid by subjects (Figure S7A). ii). The threshold should remain stable session by session (Figure S7B).

### Electrophysiology

Neural activity in M1 was recorded using The Cerebus Neural Signal Processor (Blackrock Neurotech). The recordings began 1 week before the task learning as the animal freely moved in the operant box. A pair of units that exhibited the highest recording stability and signal-to-noise ratio were assigned as “direct units” on the first day of learning. This sorting template was remained unchanged over the course of learning. Besides, all recorded neural activity would go through an offline spike sorting protocol comprised of valley-seeking automatic sorting and manual adjustment for *post hoc* analysis (Offline Sorter, Plexon).

The electrical stimulation to MFB was carried out by The CereStim neurostimulator (Blackrock Neurotech). We used a classic lever-pressing protocol to determine the appropriate stimulation parameters: Rats were trained to press a lever to obtain an artificial reward delivered by MFB stimulation in an operant box. One stimulation train comprised of 10 biphasic pulses at 100 Hz. Each pulse was a symmetric square wave with pulse width of 1 ms. The pulse amplitude was initialized as 40 *μ*A and increased incrementally in a step of every 5 *μ*A until the rate of lever pressing increased drastically. The pulse amplitude was then fixed in the following BMI-based training sessions. Rats exhibited different sensitivity to this artificial reward (65 ± 14.5 *μ*A).

### Behavioral and neural data analysis

#### Behavioral metrics

Task performance was evaluated by normalized success rate *SR_n_* and trial duration. The raw success rate *SR_raw_* was defined as the ratio of the number of successful trials to the total number of trials carried out in a session. We then defined the normalized success rate by:

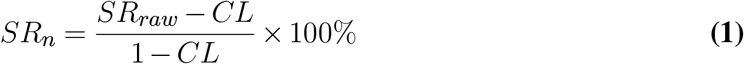

where *CL* was the chance level. Assuming the random process *NCSP*(*t*) in chance was stationary, then the chance level of the raw success rate *CL* was the average probability of exceeding the rewarding threshold in the maximum trial duration (*TD_max_* = 15*s*), denoted as *P_trial_*. We could estimate the probability density function *f*(*NCS*) by uniformly sampling NCSP in the Baseline. The corresponding percentile of the given threshold was denoted as *P_thr_* which characterized the probability of failing to exceed the threshold in one bin (0.3 s). Therefore, concerning that failing one trial follows a binomial distribution with 50 independent experiments, we have:

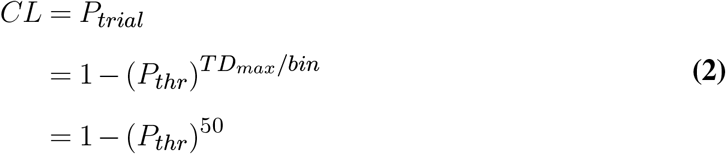

The normalization eliminated the potential noise brought by the slight drift of rewarding thresholds across sessions.

The gain of NCSP was defined as the difference of mean NCSP sampled in Task (intertrial intervals were excluded from sampling) and Baseline.

#### Data pre-processing for LFP

The raw data were first detrended in a moving window of 1 s and a stepsize of 0.5 s (Chronux, Mitra (2007)), and then low-pass filtered with 100Hz as the cutoff frequency (4^*th*^ order Butterworth zero-phase shift IIR filtering). The LFP data were extracted from the filtered data described above by down-sampling at the rate of 2000 Hz.

#### Cross-correlation histogram (CCH)

Cross-correlation histogram (CCH) is a linear statistical assessment of the interdependencies between pairs which is often used to describe functional connectivity (Vizuete et al., 2012). Raw CCHs were computed using 15 ms bin width. Specifically, the temporal offsets between any spikes from the trigger unit and all spikes from the target unit relative to that trigger spike were detected and normalized by the geometric mean of the total spikes from the trigger unit and the target unit. Hence, each bar of the CCHs reveals, for any given discharging event of the trigger unit, the probability for the target unit to fire at that particular time lead or lag. To remove the spurious correlation brought by covariation in a coarser timescale, we jittered the time of the spikes from the target unit around 15 ms (the standard deviation of the Gaussian distribution). The jittering was independently run 1000 times for each trials, then averaged to render the jittered CCH with standard errors. The corrected CCH was obtained by subtracting the jittered CCH from the raw CCH.

To statistically test the significant coincidence in particular time lead or lag, we compared the corrected CCH in given time lead or lag to its standard error.

#### Cell assembly detection

The detection algorithm was adapted from Russo and Durstewitz (2017). We limited the bin widths for analysis from 5 to 30 ms in a 5 ms step. The maximum lags were specified to 5 for each bin width, which rendered the detection range as ±25 ms to ±150 ms. We used the number of assemblies, assembly size, and precision (bin width) to characterize the fine timescale population dynamics. If two assemblies were made of the same units but differed in the temporal relationship (lags), they were seen as independent cell assemblies. The number of units contained in an assembly was referred to as assembly size. We sieved the cell assemblies that comprised of the same units in the Trial start and Pre-reward for precision comparison.

#### PCA

We used the PCA reveal the evolving patterns of population activity in a coarser timescale. Spikes were binned in 20 ms and convolved with a gaussian kernel (standard deviation = 100 ms). All successful trials were concatenated to form data matrix *D* ∈ *R*^*T*×*N*^, where *T* is the time in 20 ms bins, and *N* is the number of recorded units. PCA then decomposed *D* into scores matrix *U* ∈ *R*^*T*×*K*^ and loadings matrix *V* ∈ *R*^*N*×*K*^ such that *D* = *U* × *V^T^*, where *K* is the number of principal components (PCs). From *U*, we extracted the data in the 1 s window preceding the rewards and averaged over trials and top PCs to obtain the top PC projections. From the top PCs of one subject, we found those whose projections exhibited significant growing trends and sum them up to get the ramp-up PC of the subject. Their corresponding weights were also summed up to render the projection weights of ramp-up PC.

The dynamic ranges of ramp-up PC projection and the number of assembly is computed by subtracting the first tertile from the last tertile of the corresponding trajectories.

#### LFP-LFP coherence

We used directed coherence *γ_ij_*(*f*) to characterize directed functional connectivity from *j* to *i* (the eMVAR Matlab Toolbox, Faes and Nollo (2011)). The directed coherence is a spectral representation of a multivariate autoregressive (MVAR) model. We first fit the causal MVAR model:

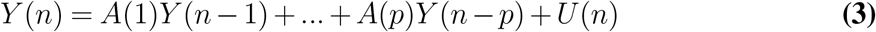

where *Y*(*n*) is a vector ℝ^2^, *A*(*k*) is a 2 × 2 coefficient matrix regressing *k*(*k* = 1,2…*p*) timesteps forward. In our case, *p* corresponded to 15 ms before the regressed time, which corresponded to the fine timescale defined throughout the study. *U*(*n*) is the estimated residuals. Least squares was used to estimate the time series. The MVAR model coefficients **A** = [*A*(1)…*A*(*p*)] was further transformed to the frequency domain as **H**(*f*). Then the directed coherence can be given by:

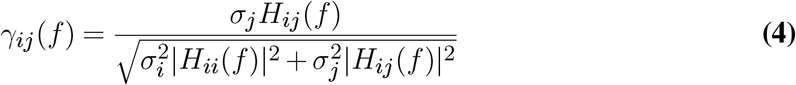

Besides, we used a model-free measure of coherence *C_ij_* to capture undirected connectivity between channel *i* and *j* (coherencyc from Chronux Matlab package, Mitra (2007)). Specifically:

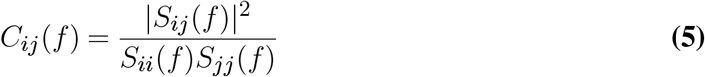

where *S_ij_*(*f*) is the cross-spectral density of the signal, and *S_ii_*(*f*) and *S_jj_*(*f*) are the power spectral density functions.

For both of the measures above, we averaged them over 7 to 30 Hz to yield coherences in the low-frequency band.

#### LFP power estimation and phase-locking

The power of theta oscillation and gamma oscillation was estimated using *mtspectrumc* in Chrounx on data from three periods: Baseline, Trial start and Pre-reward. We used the theoretical errors to pair test whether the power in the Pre-reward was significantly higher than those in the Trial start and the Baseline.

To compare the level of spike phase locking to the common average of LFP, we used the phase locking value (PLV), which has been widely used in pyramids of studies. We first inferred the phase of LFP *ϕ*(*t*) from the analytic signal *z*(*t*) via the Hilbert transform:

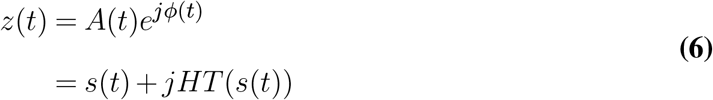

The PLV was computed by averaging the instantaneous circular phases at the time when the neuron spiked *t* = *t*_1_, *t*_2_,…, *t_k_* (Lachaux et al. (1999)):

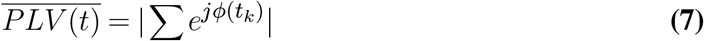

### Network simulation

#### Network model

The network consisted of *N_E_* = 4000 excitatory and *N_I_* = 1000 inhibitory leaky integrate-and-fire (LIF) neurons:

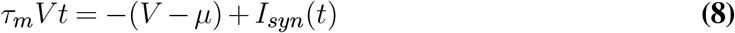

with voltage threshold for spiking *V_th_* = 1. After the spike was emitted, the neuron went through a refractory period of 5 ms and was reset to *V*_0_ = 0.

Neurons were randomly connected with probability *p^xy^*. We inherited the parameters primarily from (Litwin-Kumar and Doiron, 2012) to preserve the basic network dynamics that emerged with the clustering structure. *τ_m_* is the membrane time constant, *μ* is the bias term, *I_syn_* is the sum of synaptic inputs from other connected neurons:

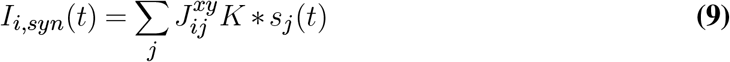

where 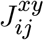 denoted the synaptic strength of the synapse from the neuron *j* in the population *y* to the neuron *i* in the population *x,x,y* ∈ *E,I*. *s_j_*(*t*) is the spike train that neuron *j* sent, *K* is the alpha function kernel:

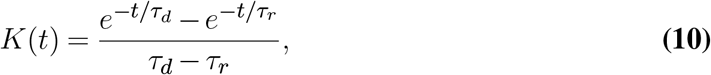

with *τ_d_* denoting the decay time and *τ_r_* denoting the rise time of the synaptic conductance dynamics. The constants for constructing basic neural network were summarized in Table 1

**Table 1.** Constants of network simulation.

| Constant | Value | Description |
| --- | --- | --- |
| $p^{EI}, p^{IE}, p^{II}$ | 0.5 | connection probability |
| $p^{EE}$ | 0.19 | connection probability for neurons both from excitatory population |
| $\tau_m^E$ | 15 ms | membrane time constant for excitatory neurons |
| $\tau_m^I$ | 10 ms | membrane time constant for inhibitory neurons |
| $\mu_E$ | 1.1 ~ 1.2 | random bias |
| $\mu_I$ | 1 ~ 1.05 | random bias |
| $J_{EE}$ | 0.024 | synaptic strength from excitatory neuron to excitatory neuron |
| $J_{IE}$ | 0.014 | synaptic strength from excitatory neuron to inhibitory neuron |
| $J_{EI}$ | -0.045 | synaptic strength from inhibitory neuron to excitatory neuron |
| $J_{II}$ | -0.056 | synaptic strength from inhibitory neuron to inhibitory neuron |
| $\tau_d^E$ | 3 ms | decay time of excitatory synapses |
| $\tau_d^I$ | 2 ms | decay time of inhibitory synapses |
| $\tau_r$ | 1 ms | rise time of synapses |

To construct a subnetwork that has stronger and denser connections, we exploited the concept of “cluster” developed in Litwin-Kumar and Doiron (2012) with *R_EE_* being modulated to render different degree of connectivity. In addition to the difference in connection probabilities, the synaptic strength of synapses within the subnetwork was stronger than other synapses. Therefore, we have the probability of connection within the subnetwork *p_EE,in_* = *R_EEpEE_* and the synapses within the subnetwork *J_EE,in_* = *R_EE_ J_EE_*.

In the learning experiment, *J_EE_* could be changed following the reward-based STDP rule for the experiment group and random reward group, and fixed for the control group. All of these groups started under identical initial conditions in the simulations. Similar to the *in vivo* session structure, the NCSP threshold for reward was set according to a 5-minute baseline recording. Once the NCSP crossed the threshold, the weights of all connections between excitatory neurons *W_ji,EE_* in the network were updated based on their eligibility traces *c_ji_*(*t*) (Legenstein et al. (2008)):

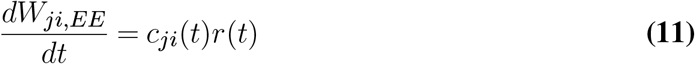

where *r*(*t*) = ∑_*n*_ *δ*(*t* – *t_n_*) is the reward signal comprised of Dirac functions at rewarding times *t*_1_, *t*_2_,…, *t_n_*, eligibility traces *c_ji_* is given by the convolution of kernel function *f_c_*(*t*) and spurious derivative of weights *w_ij,EE_*:

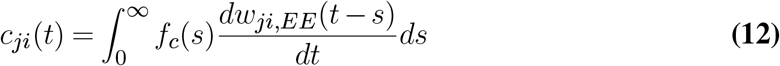

The eligibility trace kernel *f_c_*(*t*) took the forms of alpha function:

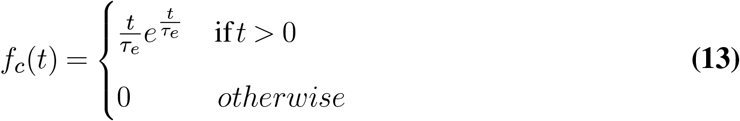

where *τ_e_* is the time constant set to 100 ms. And the spurious derivative of weights *w_ij,EE_* following the STDP rule was given by:

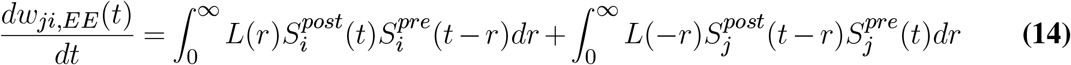

where *L*(·) was the STDP learning curve that concluded from a line of experimental researches:

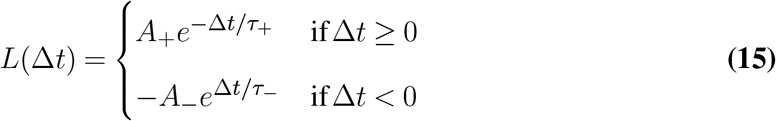

We limit the weights to the range of 0 to *W_max_* to prevent the weights from exploding.

A learning session was terminated if the number of successful trials in the experiment group was 20 times higher than the control group within the maximum duration.

For random reward group, the above STDP rule held with the rewarding time being jittered in ±3 s. Therefore, the number of times the weights were updated was identical to that of the experiment group.

The constants for the simulation of reward-based STDP learning are summarized in Table 2.

**Table 2.** Constants in reward-based STDP learning simulation.

| Constant | Value | Description |
| --- | --- | --- |
| $g$ | 12 | inhibition-excitation ratio |
| $J_{EE}$ | 0.024 | synaptic strength from excitatory neuron to excitatory neuron |
| $J_{IE}$ | 0.014 | synaptic strength from excitatory neuron to inhibitory neuron |
| $J_{EI}$ | -0.045 | synaptic strength from inhibitory neuron to excitatory neuron |
| $J_{II}$ | -0.056 | synaptic strength from inhibitory neuron to inhibitory neuron |
| $W_0^{EE}, W_0^{IE}$ | $J_{EE}, J_{IE}$ | initial weights for all excitatory connections |
| $W_0^{EI}, W_0^{II}$ | $-gJ_{EI}, -gJ_{II}$ | initial weights for all inhibitory connections |
| $W_{max}$ | $2W_0^{EE}$ | Maximum value of weights |
| $A_+$ | $0.01W_{max}$ | coefficient of positive STDP learning curve |
| $A_-$ | $1.05A_+$ | coefficient of negative STDP learning curve |
| $\tau_+, \tau_-$ | 30 ms | time constant of STDP learning curve |
| $\nu$ | 0.01 | external input to individual neurons at every simulation step |

#### Numerical simulation

The simulation was proceeded every 0.1 ms. For simulations in Figure 5 and Figure 6, each combination of *ν_ext_* and *g* was run for 30 s. For reward-based learning simulation in Figure 7, we performed 100 runs for both types of network. Each run was given a maximum simulation time of 1350 s which approximated 5-session *in vivo* learning course

#### Network metrics

Mean firing rates, coefficient of variation of inter-spike intervals, and the proportion of synchronous pairs were incorporated to characterize cluster dynamics given external stimulus *ν_ext_* and effective inhibition *g* (Kumar et al., 2008). The unit value of *ν_ext_* was *ν* shown in Table 2, thus *ν_ext_* was a multiplier. Because the conductance dynamics was a fast transient, we can simplify it as follows:

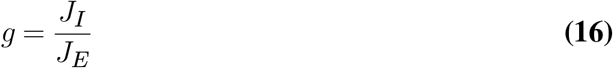

*J_I_* and *J_E_* were the synaptic efficacy from inhibitory and excitatory neurons respectively. To approximate the setting in the *in vivo* experiment, we binned the simulated spike trains in 15 ms before calculating the proportion of synchronous pairs.

The clustering coefficient of node *i* in a weighted directed network (WDN) is defined as the fraction of the geometric average of the subgraph edge weights (Fagiolo (2007)), given by:

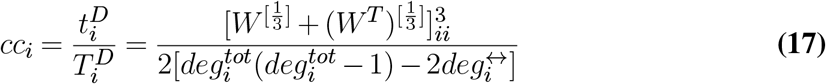

where the 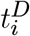 is all directed triangles derived by node *i* and 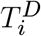 is all possible triangles that node *i* could derive. *W* = *w_ij_* is the weight matrix. 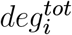 represents the total degree that includes all in-degree and out-degree of node *i*, while 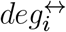 is the number of bidirectional edges to node *i*. Because Eq.17 is an extension from binary directed networks (BDNs), we scaled the weights in the weight matrix by the *W_max_* to [0, 1]

To quantify the overall level of clustering given a set of nodes, the average clustering coefficient 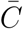 was used:

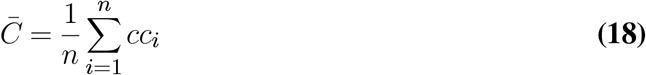

In the reward-based learning simulation, we used this measure on all nodes that are connected to the target neuron to quantify the clustering effect before and after learning.

The synchrony defined in the reward-based learning simulation (Figure 7I, S5G) is different from the one defined by the proportion of synchronous pairs. Since we already set the *ν_ext_* and *g* for network to operate on SI regime in the learning simulation, we used the average pair-wise correlation coefficient of the cluster with N neurons to define the synchrony *S* with higher sensibility:

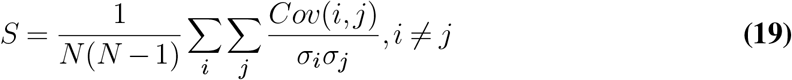

#### Network visualization

We used NetworkX for visualization. The layout was drawn using the Fruchterman-Reingold force-directed algorithm to best reveal the subnetwork structure. Connections were undirected and unweighted in Figure 6A to capture the essence of network structure.

## Data and Code Availability

Pre-processed data and codes will be available upon publication. Raw data will be available upon reasonable request at the time of publication.

## Author Contributions

Y.N. and S.Z. conceived and designed the study. Y.N.,T.Z.,G.W, and J.H. performed the animal experiment. Y.N. performed the computer simulations. Y.N. performed the analysis. Y.N. wrote the manuscript with input from T.Z.,T.L. and S.Z.. Y.N., T.L. and S.Z. revised the manuscript.

## Supplementary Information

**Figure S1.**
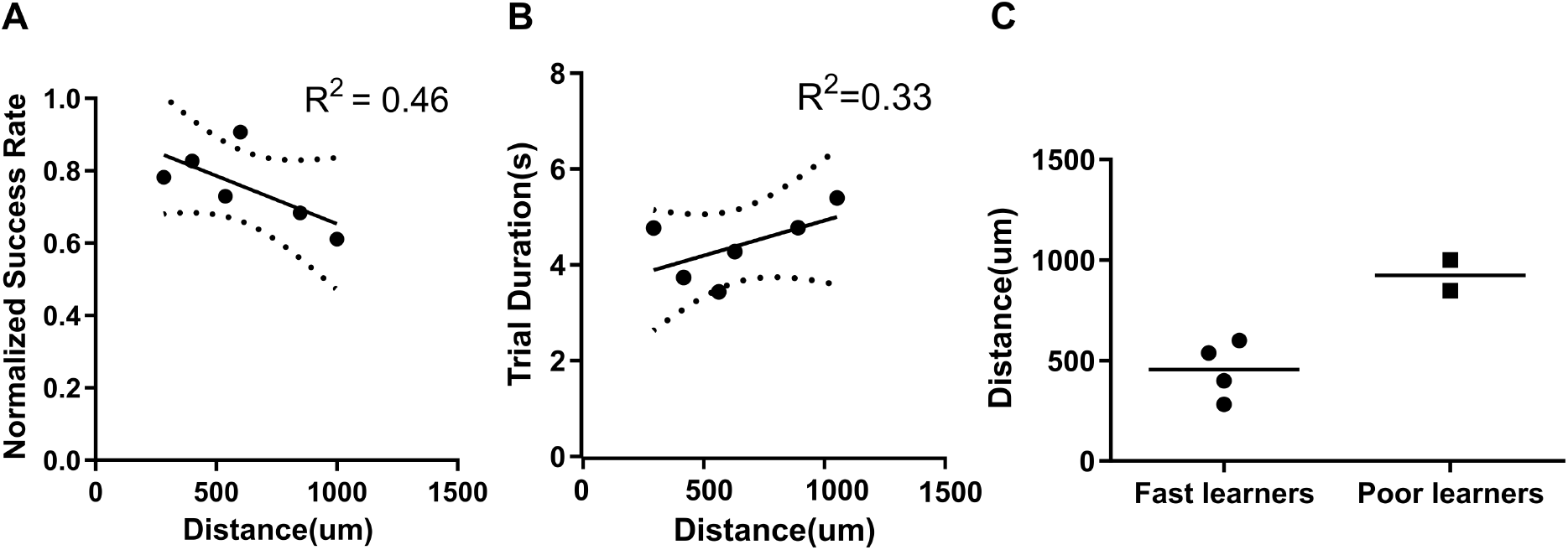
Behavioral performance varies with distance between direct neurons. (**A**) Linear regression of success rate and electrodes distance. 95% confidence band was circumvented by the dashed lines. (**B**) Linear regression of trial duration and electrodes distance. 95% confidence band was circumvented by the dashed lines. (**C**) Distance between the trigger unit and the target unit in each groups.

**Figure S2.**
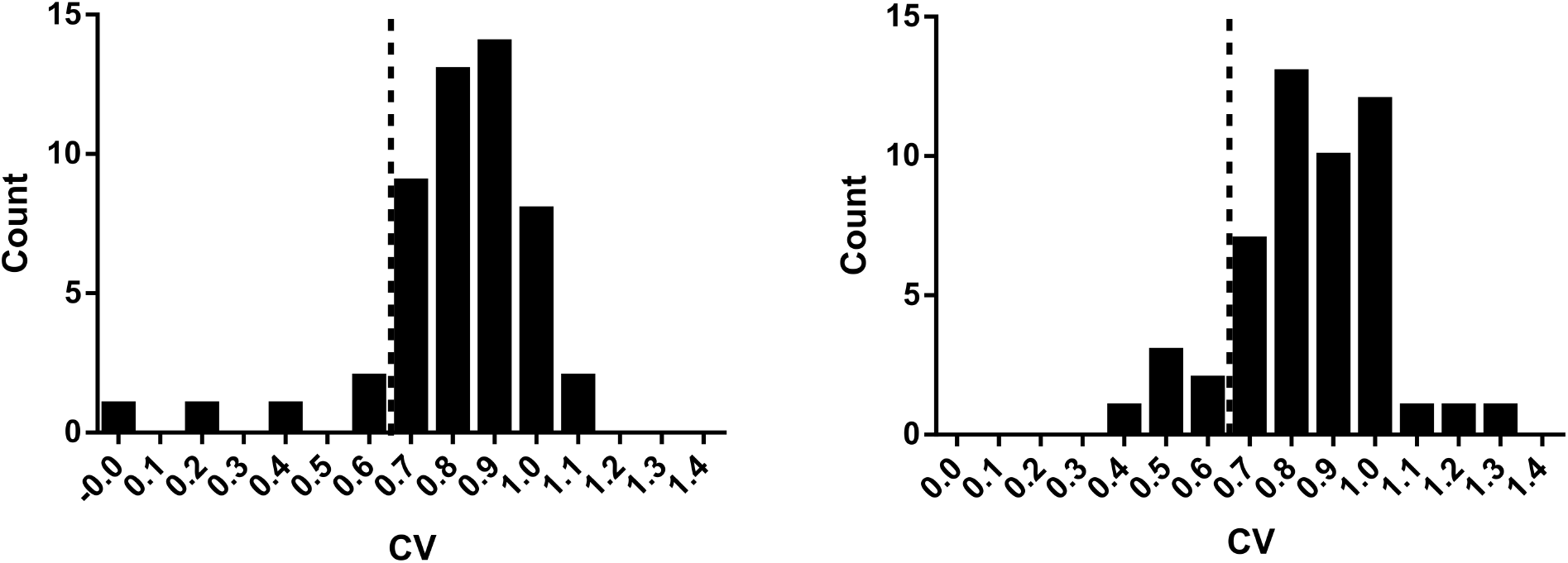
Irregular firings in Pre-reward (*left*) and Trial start (*right*).

**Figure S3.**
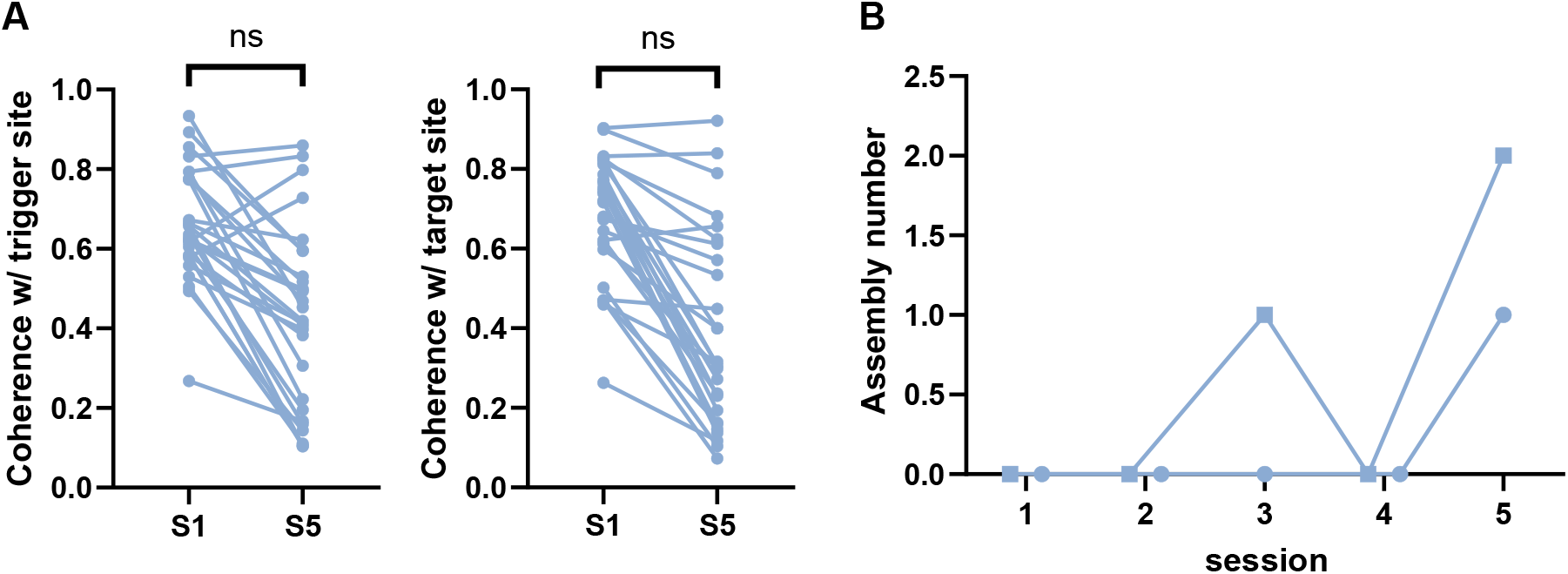
(**A**) *Left*: Coherence computed between the trigger site and other indirect sites (paired t-test, n = 28 from 2 “poor learners”). *Right*: Coherence computed between the target site and other indirect sites (paired t-test, n = 28 from 2 “poor learners”). (**B**) The evolution of assembly number in Pre-reward over the course of learning, error bars represent the SEMs. Samples are staggered. Linear regression, n = 10 from 2 “poor learners”, samples are staggered, R^2^ = 0.41, p = 0.046 for slope significantly nonzero.

**Figure S4.**
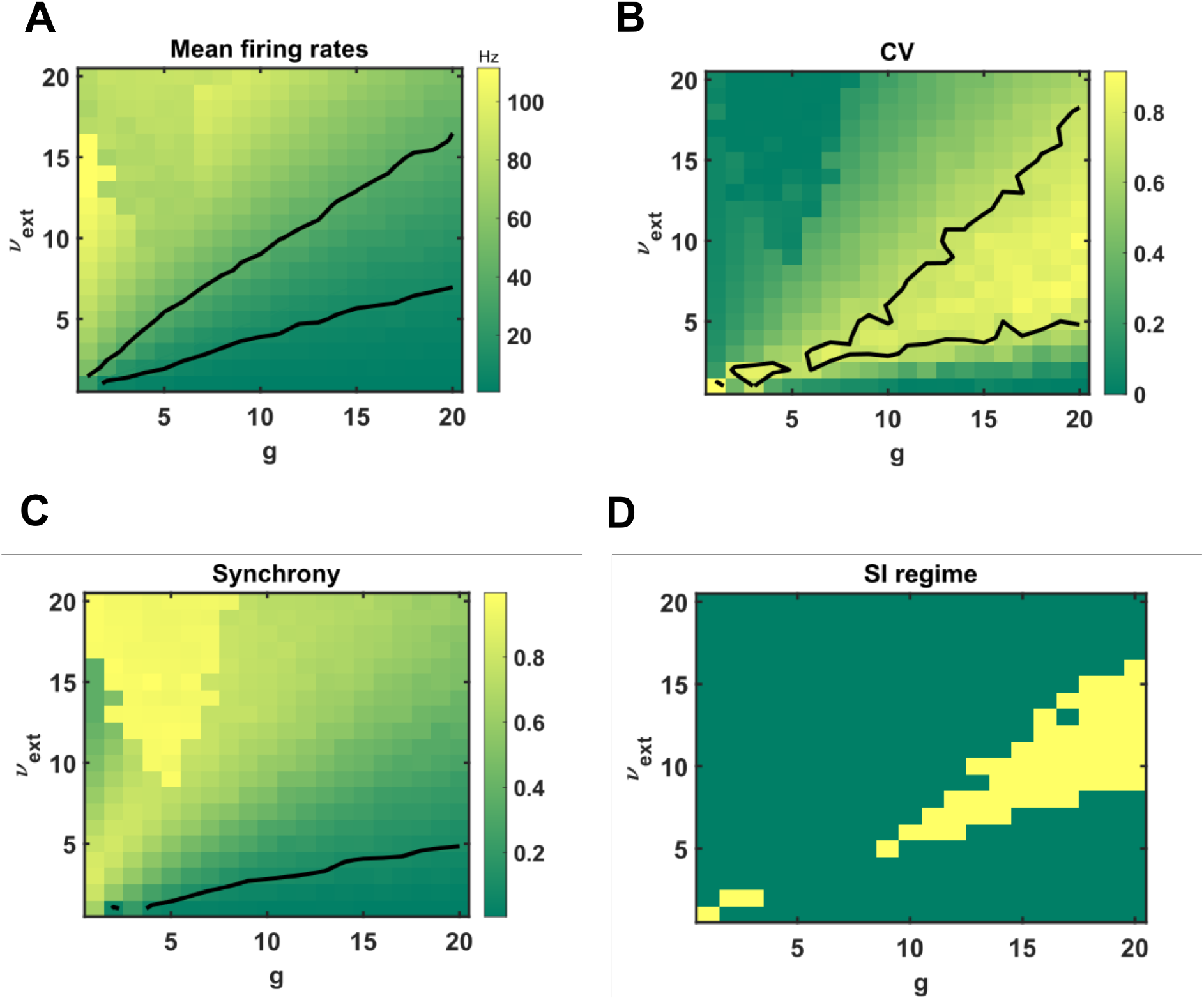
Population activity and dynamical regimes under *R_EE_*=2.5.

**Figure S5.**
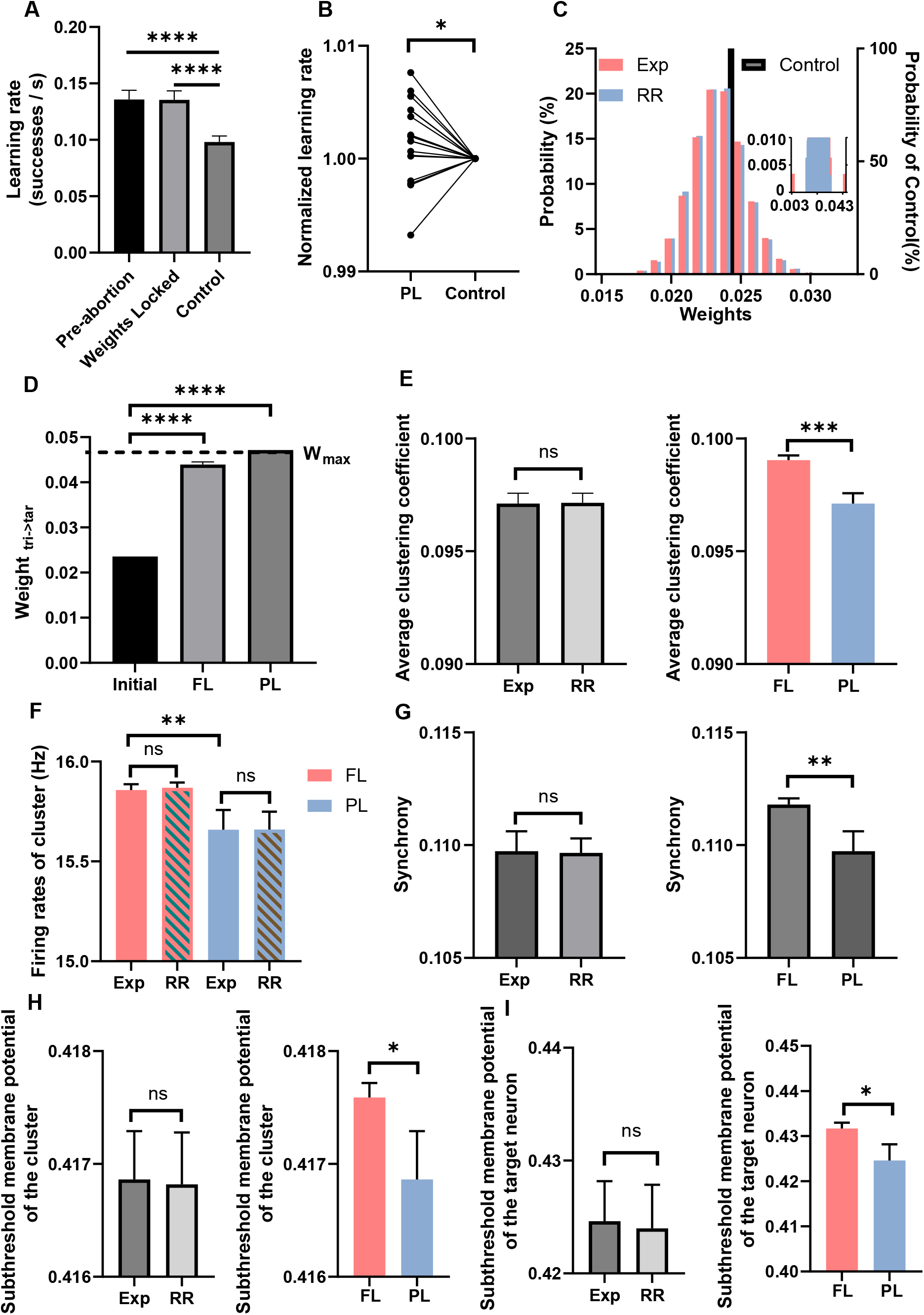
(**A**) The learning rates estimated under three conditions (n = 98, Wilcoxon matched-pairs signed rank test). (**B**) The learning rate of “poor learners” normalized by that of its control groups is greater than 1 (n = 18, paired t-test, p = 0.0357). (**C**). Example of the distribution of network weights (E-E connection only) in three groups. *Inset*: Zoomed histogram to the probability under 0.01%. (**D**) The weights of connection from the trigger neuron to the target neuron increased in both the “fast learners” and “poor learners” compared to the initial weights. (**E**) *Left*: No difference in the average clustering coefficient between the experiment group and the random reward group of “poor learners” (n = 18, paired t-test, p = 0.69, mean±SEM). *Right*: The average clustering coefficient of “fast learners” (*coral*, n = 80) is higher than that of the “poor learners” (*blue*, n = 18) (One tailed t-test, p = 0.0001, mean±SEM). (**F**) No difference in the average firing rates of cluster between the experiment group and the random reward group of both “fast learners” (*coral*, n = 80, paired t-test, p = 0.85, mean±SEM) and “poor learners” (*blue*, n = 18, paired t-test, p = 0.52, mean±SEM). But firing rates of “fast learners” were higher than that of “poor learners” (One tailed t-test, p = 0.006, mean±SEM). (**G**) *Left*: No difference in the synchrony between the experiment group and the random reward group of “poor learners” (n = 18, paired t-test, p = 0.69, mean±SEM). *Right*: The synchrony of “fast learners” (*coral*, n = 80) is higher than that of the “poor learners” (*blue*, n = 18) (One tailed t-test, p = 0.0024, mean±SEM). (**H**) *Left*: No difference in the membrane potential of cluster between the experiment group and the random reward group of “poor learners” (n = 18, paired t-test, p = 0.79, mean±SEM). *Right*: The membrane potential of cluster of “fast learners” (*coral*, n = 80) is higher than that of the “poor learners” (*blue*, n = 18) (One tailed t-test, p = 0.017, mean±SEM). (**I**) *Left*: No difference in the membrane potential of the target neuron between the experiment group and the random reward group of “poor learners” (n = 18, paired t-test, p = 0.31, mean±SEM). *Right*: The membrane potential of the target neuron of “fast learners” (*coral*, n = 80) is higher than that of the “poor learners” (*blue*, n = 18) (One tailed t-test, p = 0.013, mean±SEM).

**Figure S6.**
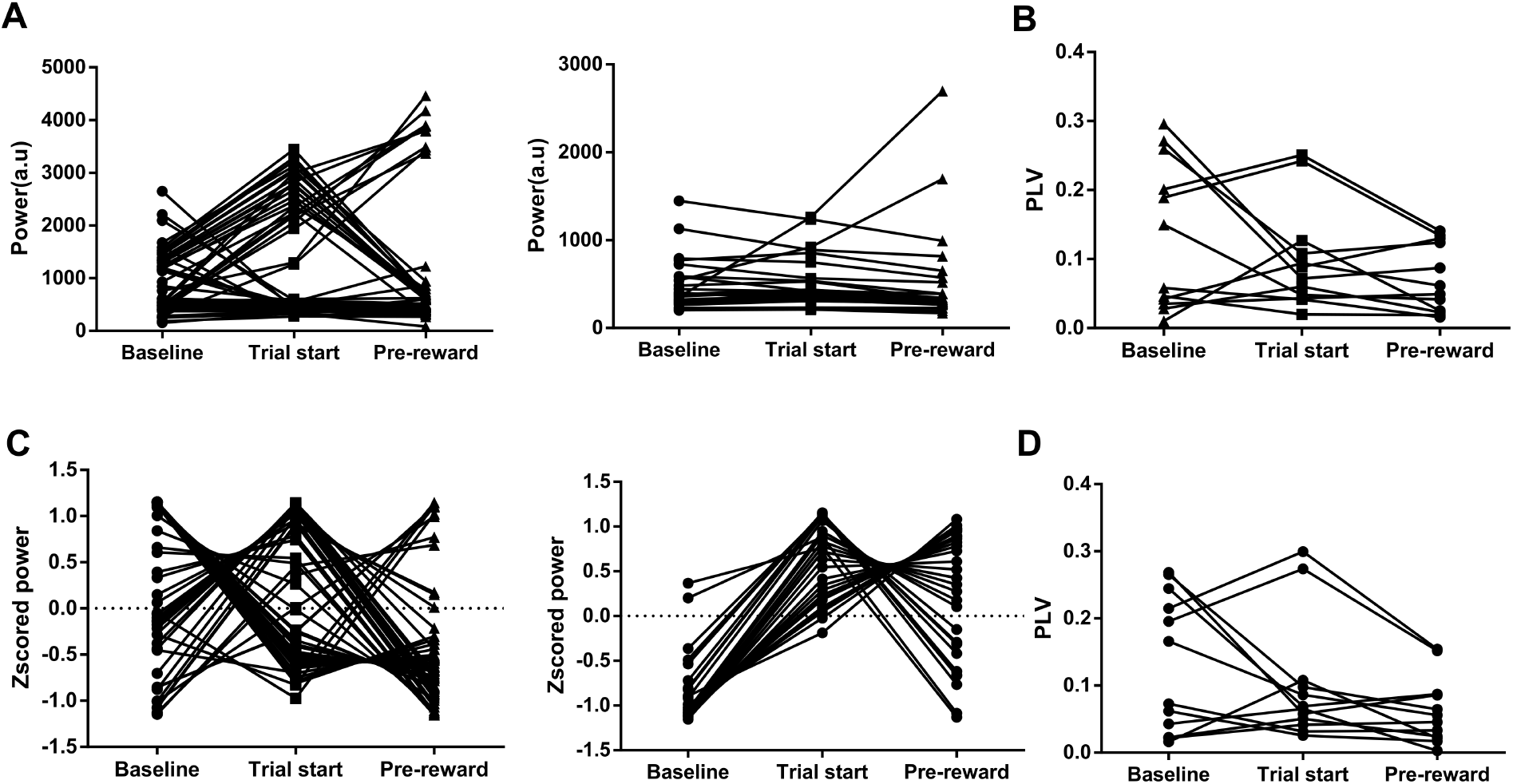
LFP power and spike phase-locking under different periods. (**A**). *Left*: Gamma band power of all channels from the “fast learners”. 15/60 significantly higher in the Pre-reward than the Trial start and the Baseline. *Right*: Gamma band power of all channels from the “slow learners”. 2/30 significantly higher in the Pre-reward than the Trial start and the Baseline. (**B**). Phase-locking values based on averaged LFP in gamma band pooling over all direct neurons and rats (n = 12). One-tailed t-test: Pre-reward < Baseline, P=0.006; Pre-reward < Trial start, p = 0.04. (**C**). *Left*: Normalized theta band power of all channels from the “fast learners”. *Right*: Normalized theta band power of all channels from the “slow learners”. (**D**). Phase-locking values based on averaged LFP in theta band pooling over all direct neurons and rats (n = 12). One-tailed t-test: Pre-reward < Baseline, p = 0.0037; Pre-reward < Trial start, P = 0.016.

**Figure S7.**
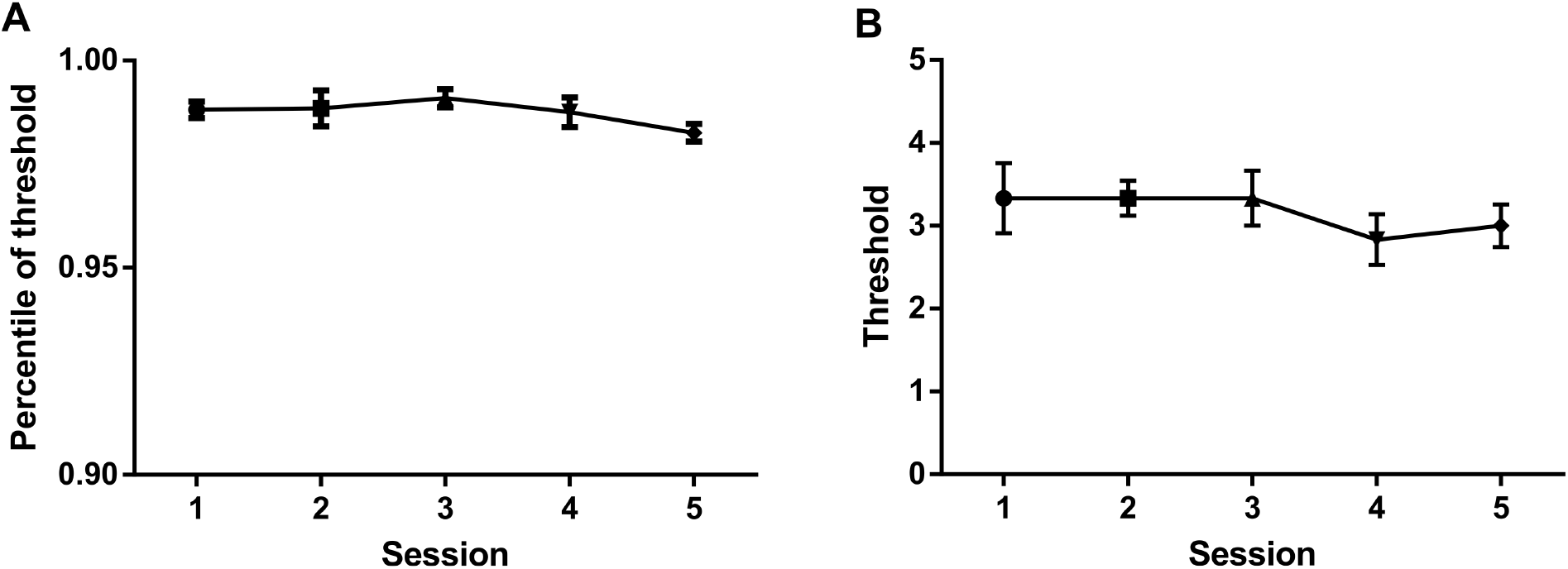
The thresholds were stable across sessions. **A**. The corresponding percentile of the threshold across sessions. mean±SEM **B**. The value of the threshold across sessions. mean±SEM

## References

Abeles, M. Role of the cortical neuron: integrator or coincidence detector? Israel journal of medical sciences, 18(1):83–92, 1982.

Abeles, M. Corticonics: Neural circuits of the cerebral cortex. Cambridge University Press, 1991.

Albert, C. Y. and Margoliash, D. Temporal hierarchical control of singing in birds. Science, 273(5283):1871–1875, 1996.

Amarasingham, A., Harrison, M. T., Hatsopoulos, N. G., and Geman, S. Conditional modeling and the jitter method of spike resampling. Journal of Neurophysiology, 107(2):517–531, 2012.

Ames, K. C. and Churchland, M. M. Motor cortex signals for each arm are mixed across hemispheres and neurons yet partitioned within the population response. Elife, 8:e46159, 2019.

Artola, A., Bröcher, S., and Singer, W. Different voltage-dependent thresholds for inducing long-term depression and long-term potentiation in slices of rat visual cortex. Nature, 347(6288):69–72, 1990.

Athalye, V. R., Carmena, J. M., and Costa, R. M. Neural reinforcement: re-entering and refining neural dynamics leading to desirable outcomes. Current opinion in neurobiology, 60:145–154, 2020.

Baker, S., Spinks, R., Jackson, A., and Lemon, R. Synchronization in monkey motor cortex during a precision grip task. i. task-dependent modulation in single-unit synchrony. Journal of Neurophysiology, 85(2):869–885, 2001.

Banerjee, A., Seriès, P., and Pouget, A. Dynamical constraints on using precise spike timing to compute in recurrent cortical networks. Neural Computation, 20(4):974–993, 2008.

Baranauskas, G. Can optogenetic tools determine the importance of temporal codes to sensory information processing in the brain? Frontiers in systems neuroscience, 9:174, 2015.

Barlow, H. B. et al. Possible principles underlying the transformation of sensory messages. Sensory communication, 1(01), 1961.

Batista, A. Brain-computer interfaces for basic neuroscience. In Handbook of clinical neurology, volume 168, pages 233–247. Elsevier, 2020.

Berger, D., Warren, D., Normann, R., Arieli, A., and Grün, S. Spatially organized spike correlation in cat visual cortex. Neurocomputing, 70(10-12):2112–2116, 2007.

Bi, G.-q. and Poo, M.-m. Synaptic modifications in cultured hippocampal neurons: dependence on spike timing, synaptic strength, and postsynaptic cell type. Journal of neuroscience, 18(24):10464–10472, 1998.

Biane, J. S., Takashima, Y., Scanziani, M., Conner, J. M., and Tuszynski, M. H. Thalamocortical projections onto behaviorally relevant neurons exhibit plasticity during adult motor learning. Neuron, 89(6):1173–1179, 2016.

Bienenstock, E. L., Cooper, L. N., and Munro, P. W. Theory for the development of neuron selectivity: orientation specificity and binocular interaction in visual cortex. Journal of Neuroscience, 2(1):32–48, 1982.

Brette, R. Philosophy of the spike: rate-based vs. spike-based theories of the brain. Frontiers in systems neuroscience, 9:151, 2015.

Brette, R. Is coding a relevant metaphor for the brain? Behavioral and Brain Sciences, 42, 2019.

Brody, C. D. Correlations without synchrony. Neural computation, 11(7):1537–1551, 1999.

Brunel, N. Dynamics of sparsely connected networks of excitatory and inhibitory spiking neurons. Journal of computational neuroscience, 8(3):183–208, 2000.

Buzsáki, G. Theta oscillations in the hippocampus. Neuron, 33(3):325–340, 2002.

Carmena, J. M., Lebedev, M. A., Crist, R. E., O’Doherty, J. E., Santucci, D. M., Dimitrov, D. F., Patil, P. G., Henriquez, C. S., Nicolelis, M. A. L., and Segev, I. Learning to control a brain–machine interface for reaching and grasping by primates. PLoS biology, 1(2):e42, 2003.

Chapin, J. K., Moxon, K. A., Markowitz, R. S., and Nicolelis, M. A. Real-time control of a robot arm using simultaneously recorded neurons in the motor cortex. Nature neuroscience, 2(7):664–670, 1999.

Cheney, P. D. and Fetz, E. E. Functional classes of primate corticomotoneuronal cells and their relation to active force. Journal of neurophysiology, 44(4):773–791, 1980.

Chi, Z. and Margoliash, D. Temporal precision and temporal drift in brain and behavior of zebra finch song. Neuron, 32(5): 899–910, 2001.

Churchland, M. M., Cunningham, J. P., Kaufman, M. T., Foster, J. D., Nuyujukian, P., Ryu, S. I., and Shenoy, K. V. Neural population dynamics during reaching. Nature, 487(7405):51–56, 2012.

Clancy, K. B. and Mrsic-Flogel, T. D. The sensory representation of causally controlled objects. Neuron, 109(4):677–689, 2021.

Clancy, K. B., Koralek, A. C., Costa, R. M., Feldman, D. E., and Carmena, J. M. Volitional modulation of optically recorded calcium signals during neuroprosthetic learning. Nature neuroscience, 17(6):807–809, 2014.

Clopath, C., Ziegler, L., Vasilaki, E., Büsing, L., and Gerstner, W. Tag-trigger-consolidation: a model of early and late long-term-potentiation and depression. PLoS computational biology, 4(12):e1000248, 2008.

Cunningham, J. P. and Byron, M. Y. Dimensionality reduction for large-scale neural recordings. Nature neuroscience, 17(11): 1500–1509, 2014.

Cunningham, J. P. and Ghahramani, Z. Linear dimensionality reduction: Survey, insights, and generalizations. The Journal of Machine Learning Research, 16(1):2859–2900, 2015.

Davies, M., Srinivasa, N., Lin, T.-H., Chinya, G., Cao, Y., Choday, S. H., Dimou, G., Joshi, P., Imam, N., Jain, S., et al. Loihi: A neuromorphic manycore processor with on-chip learning. Ieee Micro, 38(1):82–99, 2018.

Decharms, R. C. and Zador, A. Neural representation and the cortical code. Annual review of neuroscience, 23(1):613–647, 2000.

Denman, D. J. and Contreras, D. The structure of pairwise correlation in mouse primary visual cortex reveals functional organization in the absence of an orientation map. Cerebral Cortex, 24(10):2707–2720, 2014.

DiCarlo, J. J., Zoccolan, D., and Rust, N. C. How does the brain solve visual object recognition? Neuron, 73(3):415–434, 2012.

Diesmann, M., Gewaltig, M.-O., and Aertsen, A. Stable propagation of synchronous spiking in cortical neural networks. Nature, 402(6761):529–533, 1999.

Engelhard, B., Ozeri, N., Israel, Z., Bergman, H., and Vaadia, E. Inducing gamma oscillations and precise spike synchrony by operant conditioning via brain-machine interface. Neuron, 77(2):361–375, 2013.

Evarts, E. V. Relation of pyramidal tract activity to force exerted during voluntary movement. Journal of neurophysiology, 31 (1):14–27, 1968.

Faes, L. and Nollo, G. Multivariate frequency domain analysis of causal interactions in physiological time series. In Laskovski, A. N., editor, Biomedical Engineering, Trends in Electronics, chapter 21. IntechOpen, Rijeka, 2011. doi: 10.5772/13065. URL https://doi.org/10.5772/13065.

Fagiolo, G. Clustering in complex directed networks. Physical Review E, 76(2):026107, 2007.

Georgopoulos, A. P., Kalaska, J. F., Caminiti, R., and Massey, J. T. On the relations between the direction of two-dimensional arm movements and cell discharge in primate motor cortex. Journal of Neuroscience, 2(11):1527–1537, 1982.

Gerstein, G. L. and Perkel, D. H. Mutual temporal relationships among neuronal spike trains: Statistical techniques for display and analysis. Biophysical journal, 12(5):453, 1972.

Golub, M. D., Chase, S. M., Batista, A. P., and Byron, M. Y. Brain–computer interfaces for dissecting cognitive processes underlying sensorimotor control. Current opinion in neurobiology, 37:53–58, 2016.

Golub, M. D., Sadtler, P. T., Oby, E. R., Quick, K. M., Ryu, S. I., Tyler-Kabara, E. C., Batista, A. P., Chase, S. M., and Byron, M. Y. Learning by neural reassociation. Nature neuroscience, 21(4):607–616, 2018.

Gray, C. M., König, P., Engel, A. K., and Singer, W. Oscillatory responses in cat visual cortex exhibit inter-columnar synchronization which reflects global stimulus properties. Nature, 338(6213):334–337, 1989.

Guo, L., Xiong, H., Kim, J.-I., Wu, Y.-W., Lalchandani, R. R., Cui, Y., Shu, Y., Xu, T., and Ding, J. B. Dynamic rewiring of neural circuits in the motor cortex in mouse models of parkinson’s disease. Nature neuroscience, 18(9):1299–1309, 2015.

Hansel, D. and Sompolinsky, H. Chaos and synchrony in a model of a hypercolumn in visual cortex. Journal of computational neuroscience, 3(1):7–34, 1996.

Hasegawa, R., Ebina, T., Tanaka, Y. R., Kobayashi, K., and Matsuzaki, M. Structural dynamics and stability of corticocortical and thalamocortical axon terminals during motor learning. PloS one, 15(6):e0234930, 2020.

Hatsopoulos, N. G., Ojakangas, C. L., Paninski, L., and Donoghue, J. P. Information about movement direction obtained from synchronous activity of motor cortical neurons. Proceedings of the National Academy of Sciences, 95(26):15706–15711, 1998.

Hatsopoulos, N. G. Columnar organization in the motor cortex. Cortex; a journal devoted to the study of the nervous system and behavior, 46(2):270, 2010.

Hayashi-Takagi, A., Yagishita, S., Nakamura, M., Shirai, F., Wu, Y. I., Loshbaugh, A. L., Kuhlman, B., Hahn, K. M., and Kasai, H. Labelling and optical erasure of synaptic memory traces in the motor cortex. Nature, 525(7569):333–338, 2015.

Hennig, J. A., Golub, M. D., Lund, P. J., Sadtler, P. T., Oby, E. R., Quick, K. M., Ryu, S. I., Tyler-Kabara, E. C., Batista, A. P., Byron, M. Y., et al. Constraints on neural redundancy. Elife, 7:e36774, 2018.

Hernandez, G., Hamdani, S., Rajabi, H., Conover, K., Stewart, J., Arvanitogiannis, A., and Shizgal, P. Prolonged rewarding stimulation of the rat medial forebrain bundle: neurochemical and behavioral consequences. Behavioral neuroscience, 120 (4):888, 2006.

Hochberg, L. R., Serruya, M. D., Friehs, G. M., Mukand, J. A., Saleh, M., Caplan, A. H., Branner, A., Chen, D., Penn, R. D., and Donoghue, J. P. Neuronal ensemble control of prosthetic devices by a human with tetraplegia. Nature, 442(7099): 164–171, 2006.

Hopfield, J. J. Neural networks and physical systems with emergent collective computational abilities. Proceedings of the national academy of sciences, 79(8):2554–2558, 1982.

Hosp, J. A., Pekanovic, A., Rioult-Pedotti, M. S., and Luft, A. R. Dopaminergic projections from midbrain to primary motor cortex mediate motor skill learning. Journal of Neuroscience, 31(7):2481–2487, 2011.

Huxter, J., Burgess, N., and O’Keefe, J. Independent rate and temporal coding in hippocampal pyramidal cells. Nature, 425 (6960):828–832, 2003.

Ikegaya, Y., Aaron, G., Cossart, R., Aronov, D., Lampl, I., Ferster, D., and Yuste, R. Synfire chains and cortical songs: temporal modules of cortical activity. Science, 304(5670):559–564, 2004.

Jackson, A., Mavoori, J., and Fetz, E. E. Long-term motor cortex plasticity induced by an electronic neural implant. Nature, 444(7115):56–60, 2006.

Jadhav, S. P., Wolfe, J., and Feldman, D. E. Sparse temporal coding of elementary tactile features during active whisker sensation. Nature neuroscience, 12(6):792–800, 2009.

Jazayeri, M. and Movshon, J. A. Optimal representation of sensory information by neural populations. Nature neuroscience, 9 (5):690–696, 2006.

Kaufman, M. T., Churchland, M. M., Ryu, S. I., and Shenoy, K. V. Cortical activity in the null space: permitting preparation without movement. Nature neuroscience, 17(3):440–448, 2014.

Kleim, J. A., Hogg, T. M., VandenBerg, P. M., Cooper, N. R., Bruneau, R., and Remple, M. Cortical synaptogenesis and motor map reorganization occur during late, but not early, phase of motor skill learning. Journal of Neuroscience, 24(3):628–633, 2004.

Kobayashi, R., Kurita, S., Kurth, A., Kitano, K., Mizuseki, K., Diesmann, M., Richmond, B. J., and Shinomoto, S. Reconstructing neuronal circuitry from parallel spike trains. Nature communications, 10(1):1–13, 2019.

Koralek, A. C., Jin, X., Long II, J. D., Costa, R. M., and Carmena, J. M. Corticostriatal plasticity is necessary for learning intentional neuroprosthetic skills. Nature, 483(7389):331–335, 2012.

Kumar, A., Schrader, S., Aertsen, A., and Rotter, S. The high-conductance state of cortical networks. Neural computation, 20 (1):1–43, 2008.

Kupferschmidt, D. A., Juczewski, K., Cui, G., Johnson, K. A., and Lovinger, D. M. Parallel, but dissociable, processing in discrete corticostriatal inputs encodes skill learning. Neuron, 96(2):476–489, 2017.

Lachaux, J.-P., Rodriguez, E., Martinerie, J., and Varela, F. J. Measuring phase synchrony in brain signals. Human brain mapping, 8(4):194–208, 1999.

Legenstein, R., Pecevski, D., and Maass, W. A learning theory for reward-modulated spike-timing-dependent plasticity with application to biofeedback. PLoS computational biology, 4(10):e1000180, 2008.

Leonardo, A. and Fee, M. S. Ensemble coding of vocal control in birdsong. Journal of Neuroscience, 25(3):652–661, 2005.

Li, N., Chen, S., Guo, Z. V., Chen, H., Huo, Y., Inagaki, H. K., Chen, G., Davis, C., Hansel, D., Guo, C., et al. Spatiotemporal constraints on optogenetic inactivation in cortical circuits. Elife, 8:e48622, 2019.

Litwin-Kumar, A. and Doiron, B. Slow dynamics and high variability in balanced cortical networks with clustered connections. Nature neuroscience, 15(11):1498–1505, 2012.

London, M., Roth, A., Beeren, L., Häusser, M., and Latham, P. E. Sensitivity to perturbations in vivo implies high noise and suggests rate coding in cortex. Nature, 466(7302):123–127, 2010.

Long, M. A., Jin, D. Z., and Fee, M. S. Support for a synaptic chain model of neuronal sequence generation. Nature, 468 (7322):394–399, 2010.

Lubenov, E. V. and Siapas, A. G. Hippocampal theta oscillations are travelling waves. Nature, 459(7246):534–539, 2009.

Maynard, E., Hatsopoulos, N., Ojakangas, C., Acuna, B., Sanes, J., Normann, R., and Donoghue, J. Neuronal interactions improve cortical population coding of movement direction. Journal of Neuroscience, 19(18):8083–8093, 1999.

Michaels, J. A., Dann, B., and Scherberger, H. Neural population dynamics during reaching are better explained by a dynamical system than representational tuning. PLoS computational biology, 12(11):e1005175, 2016.

Mitani, A., Dong, M., and Komiyama, T. Brain-computer interface with inhibitory neurons reveals subtype-specific strategies. Current Biology, 28(1):77–83, 2018.

Mitra, P. Observed brain dynamics. Oxford University Press, 2007.

Moran, D. W. and Schwartz, A. B. Motor cortical representation of speed and direction during reaching. Journal of neurophysiology, 82(5):2676–2692, 1999.

Moritz, C. T. Now is the critical time for engineered neuroplasticity. Neurotherapeutics, 15(3):628–634, 2018.

Muñoz-Castañeda, R., Zingg, B., Matho, K. S., Chen, X., Wang, Q., Foster, N. N., Li, A., Narasimhan, A., Hirokawa, K. E., Huo, B., et al. Cellular anatomy of the mouse primary motor cortex. Nature, 598(7879):159–166, 2021.

Murthy, V. N. and Fetz, E. E. Synchronization of neurons during local field potential oscillations in sensorimotor cortex of awake monkeys. Journal of neurophysiology, 76(6):3968–3982, 1996.

Neely, R. M., Koralek, A. C., Athalye, V. R., Costa, R. M., and Carmena, J. M. Volitional modulation of primary visual cortex activity requires the basal ganglia. Neuron, 97(6):1356–1368, 2018.

Ngezahayo, A., Schachner, M., and Artola, A. Synaptic activity modulates the induction of bidirectional synaptic changes in adult mouse hippocampus. Journal of Neuroscience, 20(7):2451–2458, 2000.

Nicolelis, M. A. Actions from thoughts. Nature, 409(6818):403–407, 2001.

Olshausen, B. A. and Field, D. J. Sparse coding of sensory inputs. Current opinion in neurobiology, 14(4):481–487, 2004.

Pandarinath, C., O’Shea, D. J., Collins, J., Jozefowicz, R., Stavisky, S. D., Kao, J. C., Trautmann, E. M., Kaufman, M. T., Ryu, S. I., Hochberg, L. R., et al. Inferring single-trial neural population dynamics using sequential auto-encoders. Nature methods, 15(10):805–815, 2018.

Patel, K., Katz, C. N., Kalia, S. K., Popovic, M. R., and Valiante, T. A. Volitional control of individual neurons in the human brain. Brain, 144(12):3651–3663, 2021.

Perich, M. G., Gallego, J. A., and Miller, L. E. A neural population mechanism for rapid learning. Neuron, 100(4):964–976, 2018.

Perkel, D. H. and Bullock, T. H. Neural coding. Neurosciences Research Program Bulletin, 1968.

Pfeiffer, M. and Pfeil, T. Deep learning with spiking neurons: opportunities and challenges. Frontiers in neuroscience, 12:774, 2018.

Rehn, M. and Sommer, F. T. A network that uses few active neurones to code visual input predicts the diverse shapes of cortical receptive fields. Journal of computational neuroscience, 22(2):135–146, 2007.

Riehle, A. and Requin, J. Monkey primary motor and premotor cortex: single-cell activity related to prior information about direction and extent of an intended movement. Journal of neurophysiology, 61(3):534–549, 1989.

Riehle, A., Grün, S., Diesmann, M., and Aertsen, A. Spike synchronization and rate modulation differentially involved in motor cortical function. Science, 278(5345):1950–1953, 1997.

Rieke, F., Warland, D., Van Steveninck, R. d. R., and Bialek, W. Spikes: exploring the neural code. MIT press, 1999.

Rioult-Pedotti, M.-S., Friedman, D., Hess, G., and Donoghue, J. P. Strengthening of horizontal cortical connections following skill learning. Nature neuroscience, 1(3):230–234, 1998.

Russo, E. and Durstewitz, D. Cell assemblies at multiple time scales with arbitrary lag constellations. Elife, 6:e19428, 2017.

Russo, A. A., Bittner, S. R., Perkins, S. M., Seely, J. S., London, B. M., Lara, A. H., Miri, A., Marshall, N. J., Kohn, A., Jessell, T. M., et al. Motor cortex embeds muscle-like commands in an untangled population response. Neuron, 97(4):953–966, 2018.

Sadtler, P. T., Quick, K. M., Golub, M. D., Chase, S. M., Ryu, S. I., Tyler-Kabara, E. C., Byron, M. Y., and Batista, A. P. Neural constraints on learning. Nature, 512(7515):423–426, 2014.

Sakellaridi, S., Christopoulos, V. N., Aflalo, T., Pejsa, K. W., Rosario, E. R., Ouellette, D., Pouratian, N., and Andersen, R. A. Intrinsic variable learning for brain-machine interface control by human anterior intraparietal cortex. Neuron, 102(3): 694–705, 2019.

Santhanam, G., Ryu, S. I., Byron, M. Y., Afshar, A., and Shenoy, K. V. A high-performance brain–computer interface. nature, 442(7099):195–198, 2006.

Sergio, L. E., Hamel-Pâquet, C., and Kalaska, J. F. Motor cortex neural correlates of output kinematics and kinetics during isometric-force and arm-reaching tasks. Journal of neurophysiology, 94(4):2353–2378, 2005.

Shadlen, M. N. and Newsome, W. T. The variable discharge of cortical neurons: implications for connectivity, computation, and information coding. Journal of neuroscience, 18(10):3870–3896, 1998.

Shen, W., Flajolet, M., Greengard, P., and Surmeier, D. J. Dichotomous dopaminergic control of striatal synaptic plasticity. Science, 321(5890):848–851, 2008.

Shenoy, K. V., Sahani, M., and Churchland, M. M. Cortical control of arm movements: a dynamical systems perspective. Annual review of neuroscience, 36:337–359, 2013.

Singer, W. Neuronal synchrony: a versatile code for the definition of relations? Neuron, 24(1):49–65, 1999.

Sjöström, P. J., Turrigiano, G. G., and Nelson, S. B. Rate, timing, and cooperativity jointly determine cortical synaptic plasticity. Neuron, 32(6):1149–1164, 2001.

Softky, W. R. and Koch, C. The highly irregular firing of cortical cells is inconsistent with temporal integration of random epsps. Journal of neuroscience, 13(1):334–350, 1993.

Softky, W. R. Irregularity in the cortical spike code: Noise or information? California Institute of Technology, 1993.

Srinivasan, R., Winter, W. R., Ding, J., and Nunez, P. L. Eeg and meg coherence: measures of functional connectivity at distinct spatial scales of neocortical dynamics. Journal of neuroscience methods, 166(1):41–52, 2007.

Stöckl, C. and Maass, W. Optimized spiking neurons can classify images with high accuracy through temporal coding with two spikes. Nature Machine Intelligence, 3(3):230–238, 2021.

Taherkhani, A., Belatreche, A., Li, Y., Cosma, G., Maguire, L. P., and McGinnity, T. M. A review of learning in biologically plausible spiking neural networks. Neural Networks, 122:253–272, 2020.

Tang, C., Chehayeb, D., Srivastava, K., Nemenman, I., and Sober, S. J. Millisecond-scale motor encoding in a cortical vocal area. PLoS biology, 12(12):e1002018, 2014.

Torre, E., Quaglio, P., Denker, M., Brochier, T., Riehle, A., and Grün, S. Synchronous spike patterns in macaque motor cortex during an instructed-delay reach-to-grasp task. Journal of Neuroscience, 36(32):8329–8340, 2016.

Treisman, M. Temporal discrimination and the indifference interval: Implications for a model of the” internal clock”. Psychological Monographs: General and Applied, 77(13):1, 1963.

van Bergen, R. S. and Kriegeskorte, N. Going in circles is the way forward: the role of recurrence in visual inference. Current Opinion in Neurobiology, 65:176–193, 2020.

Vizuete, J. A., Pillay, S., Diba, K., Ropella, K. M., and Hudetz, A. G. Monosynaptic functional connectivity in cerebral cortex during wakefulness and under graded levels of anesthesia. Frontiers in Integrative Neuroscience, 6:90, 2012.

Yamawaki, N. and Shepherd, G. M. Synaptic circuit organization of motor corticothalamic neurons. Journal of Neuroscience, 35(5):2293–2307, 2015.

